# Profiling Dynamic Patterns of Single-cell Motility

**DOI:** 10.1101/2022.09.21.508955

**Authors:** Debonil Maity, Nikita Sivakumar, Pratik Kamat, Nahuel Zamponi, Chanhong Min, Wenxuan Du, Hasini Jayatilaka, Adrian Johnston, Bartholomew Starich, Anshika Agrawal, Deanna Riley, Leandro Venturutti, Ari Melnick, Leandro Cerchietti, Jeremy Walston, Jude M. Phillip

## Abstract

Cell motility plays an essential role in many biological processes as cells move and interact within their local microenvironments. Current methods for quantifying cell motility typically involve tracking individual cells over time, but the results are often presented as averaged values across cell populations. While informative, these ensemble approaches have limitations in assessing cellular heterogeneity and identifying generalizable patterns of single-cell behaviors, at baseline and in response to perturbations. In this study, we introduce CaMI, a computational framework designed to leverage the single-cell nature of motility data. CaMI identifies and classifies distinct spatio-temporal behaviors of individual cells, enabling robust classification of single-cell motility patterns in a large dataset (n=74,253 cells). This framework allows quantification of spatial and temporal heterogeneities, determination of single-cell motility behaviors across various biological conditions, and provides a visualization scheme for direct interpretation of dynamic cell behaviors. Importantly, CaMI reveals insights that conventional cell motility analyses may overlook, showcasing its utility in uncovering robust biological insights. Together, we present a multivariate framework to classify emergent patterns of single-cell motility, emphasizing the critical role of cellular heterogeneity in shaping cell behaviors across populations.

**Teaser:** A computational framework to identify and classify single-cell motility patterns and phenotypic heterogeneity across biological conditions.

## INTRODUCTION

The dynamic movement of cells, *i.e*., cell motility, underlies critical biological processes that regulate homeostasis and disease, including immune locomotion and surveillance, and the metastasis of cancer cells from primary tumor sites^1–4^. While cell motility involves a complex coordination of cellular displacements, tortuous paths, and effective diffusivities, the current paradigm for analyzing and reporting cell motility data is often limited to individual motility parameters such as average speeds and displacements. While these approaches are informative, they offer limited capacity to assess cellular heterogeneity, extract single-cell motility patterns, and assess the abundance of sub-populations of cells that may be enriched or depleted in response to perturbations. Furthermore, single-cell profiling is critical to properly understand and interpret cellular heterogeneity of biological phenotypes.

Recent advances in single-cell technologies and analyses have enabled the detailed quantitative description of cell types, cell states, and cell fates based on genetic, epigenetic, or protein signatures^5,6^. Building on these recent advances, we rationalized that combining single-cell profiling of cell motility with modern data science approaches could offer new solutions to define functional dynamic cell behaviors and cellular heterogeneities across multiple conditions^7–11^. Furthermore, performing these single-cell behavior analyses in biological systems with known underlying molecular information (*i.e*., knockdowns of particular genes, drug treatments) presents the potential to trace the influence of specific molecular programs on dynamic cell behaviors. Despite this promise, robust quantification of cell motility patterns to inform functional behaviors and heterogeneities of cells remain challenging. This challenge is partly due to the lack of universal standards for acquiring cell motility data (*i.e*., image acquisition settings, such as total imaging duration and acquisition intervals), that lack of large publicly available cell motility datasets for comparative interrogation of cell motility patterns, and limited single-cell computational tools designed specifically for cell motility datasets.

To address this gap, we present the computational framework CaMI, which enables the harmonization of motility datasets from multiple sources having different acquisition settings and robust identification of single-cell motility patterns (**Figure 1A**). Using CaMI, we combined non-uniform mammalian cell motility datasets from *in vitro* experiments to profile 74,253 single cells spanning 409 biological conditions^12–16^ (**Supplementary Table 1**) to derive emergent cell motility patterns at single-cell resolution for both short (2.5 hours) and long (8 hours) durations.

**Figure 1.**
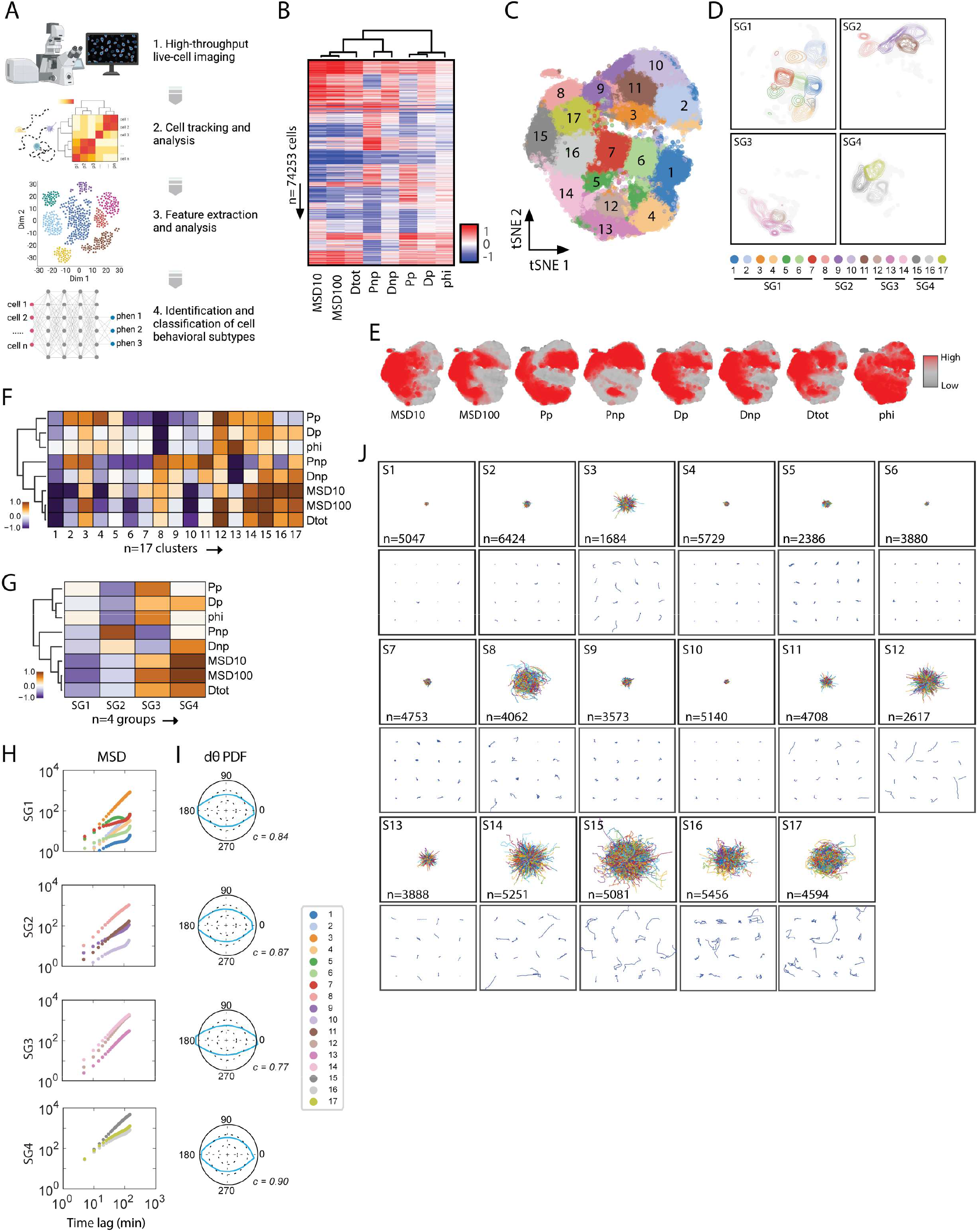
Identifying and classifying cell motility patterns across short durations. **A**. Illustration of CaMI workflow. **B**. Heatmap showing the magnitude of eight motility parameters per cell (n=74,253 cells). Each row represents a single cell, and each column represents a motility parameter. **C-D**. Two-dimensional tSNE plot showing the 17 motility clusters (**C**) where each dot is a single cell and (D) contour plots that highlight the spatial distribution of cells/clusters that are further separated into four short-duration behavior classes within the reduced dimensionality space. **E**. Two-dimensional t-SNE plot showing the magnitude of the eight motility parameters per cluster region. **F-G**. Heatmap showing the average magnitude of the eight cell motility parameters among each of the 17 cell motility clusters and the four short duration behavioral classes (G). **H**. Mean squared displacements per long duration (F) motility clusters separated based on the four short-duration behavior classes (SG1-SG4) **I**. Probability density function profiles of the angular velocity magnitudes for the four short duration behavior classes. High circularity values (less ellipsoidal profiles) indicate that cells have a similar likelihood of moving in multiple directions. **J**. Visualization of single-cell trajectories per cluster, the top panel shows the origin-centered trajectories for all cells within a given cluster, bottom panel shows 16 randomly selected cells per cluster. Scalebar denotes 100μm.

To demonstrate the biological utility of this framework, we applied CaMI to study how molecular and environmental perturbations induce fractional redistributions in cell motility behaviors in three test cases: 1) to quantify how environmental perturbations, such as cell density, can impact emergent single-cell motility properties; 2) to quantify how molecular modulators of cytoskeletal elements that are known to mechanistically impact cell movement drive distinct cell motility patterns, and 3) to quantify emergent cell motility patterns in primary human cells to profile heterogeneity in actual patients. Lastly, to demonstrate the technical utility of this framework, we show how spatial and temporal heterogeneities (computed as the entropies) across distinct motility patterns can predict the dimensionality of cells moving in either 2D or 3D microenvironments. The curated motility dataset (n=409 biological conditions) and the MATLAB code for the CaMI pipeline are available within the supplementary documents and publicly available here: https://github.com/PhillipLab-JHU/CaMI.

## RESULTS

### Description of data acquisition and curation

To develop the computational framework to identify and classify emergent patterns of single-cell motility across multiple conditions, we combined existing and newly acquired cell motility datasets^7,17–22^. Using commercial cell tracking software (Metamorph), we tracked the positions (*x, y, t*) of single cells to generate single-cell trajectories for cells across all experimental conditions presented. All cell trajectories used, and presented in this work, were derived from time-lapse movies collected using brightfield of phase-contrast images. Movies ranged in duration from 2.5 to 16 hours, with interval times and temporal resolutions ranging from 2 to 10 minutes. Below, we discuss the harmonization of the temporal resolution from these videos and the quantification of patterns within short-duration (2.5 hour) and long-duration (8 hour) videos. To derive short-duration trajectories, videos of the same cell were cut into several trajectories.

### Imaging interval times determine the shape of cell movement trajectories

Currently, there are no universal standards for acquiring cell motility data with regards to the total imaging duration or image acquisition intervals. While each researcher choses these imaging parameters based on a set of user-dependent optimization parameters (*e.g*., convenience), this often presents challenges for downstream data integration across datasets generated across multiple experiments or multiple labs. To combine motility datasets covering multiple biological conditions and from multiple sources, we first needed to integrate the data without introducing significant noise and artifacts into the datasets. We first restricted the total imaging duration to 2.5 hours. This timescale was chosen not only to maximize the total number of cells within our dataset, but also to have a timescale that is achievable for both *in vitro* and *in vivo* imaging experiments. After curating the 2.5 hr. trajectories per cell, we evaluated the effective contributions of image acquisition intervals (*i.e*., time between subsequent images—temporal resolution) on the magnitude of cell motility.

Using a test case of ht1080 fibrosarcoma cells moving within 3D Collagen-I gels, we *in-silico* varied the interval times from the most to least time-resolved trajectories within our dataset, *i.e*., 2 minutes to 10 minutes (sample trajectories shown in **Supplementary Figure 1A**, top). We varied the acquisition time by excluding intervening points from the fully time-resolved trajectory. Results indicated that the interval times significantly impacted the shapes of the cell trajectories and their corresponding motility parameters. The same trajectory acquired across different interval times had significantly different geometric concordance (**Supplementary Figure 1A**, bottom), or the average area of triangles formed between intervening points along a given trajectory (see **Methods**). Together, these results highlight that acquisition intervals contribute significantly to the magnitude of the perceived motility and that temporal resolution matters, *i.e*., cell movements based on 2-minute intervals do not equal cell movements at 10-minute intervals.

### Harmonizing single-cell motility data across multiple datasets

Since the interval times determine the shape and magnitude of the cell trajectories, to accurately compare trajectories acquired at different interval times we needed a robust approach to estimate the intervening x-y coordinates in lower resolution trajectories to harmonize the temporal resolutions of all trajectories within our dataset. Using linear interpolation, we identified spatially weighted cell locations for symmetric (8-minute to 4-minute, *i.e*., midpoint) and asymmetric (8-minute to 6-minute) multiples of the acquisition-interval times (**Supplementary Figure 1B-C**). We observed that interpolated cell locations showed significant deviations in concordance from the original trajectories in the both the symmetric and asymmetric cases (**Supplementary Figure 1B-C**). To account for this incongruency, we developed a pseudo-Monte Carlo (pMC) approach that applies a random perturbation defined by a pMC factor to the linearly interpolated locations, thereby estimating the location of cells at a given time point (see **Methods**). We found that this pMC approach generated concordances with no significant difference to the trajectories generated using actual data points for both the symmetric and asymmetric conditions (**Supplementary Figure 1B-C**, bottom).

In addition to the geometric concordance, we tested how higher order motility parameters varied based on interpolation approach. We found that the pMC approach could replicate short and long time-scale MSD behaviors of the original trajectory, while linear interpolation showed deviations at shorter time scales (**Supplementary Figure 2A**). To further compare the motility properties between actual, interpolated, and pMC-estimated trajectories we computed motility parameters using the anisotropic persistent random walk (APRW) model, which corrects for the fact that cells do not always follow random walks on 2D or 3D substrates^19,23^. Results indicated no statistical differences between actual and reconstructed trajectories using an optimized pMC-2.01 (referred to as pMC-2 moving forward) for any of the eight motility parameters assessed (**Supplementary Figure 2B**). While the pMC-2 reconstructed trajectories do not exactly map the actual/ground truth trajectories, this approach maximizes the geometric concordance, thereby recapitulating the magnitude of the higher order motility parameters. Therefore, we constructed a harmonized dataset for the 409 biological conditions with a total duration of 2.5 hours and acquisition time intervals of 5 minutes (**Supplementary Table 1**). We selected an acquisition time interval of 5 minutes because this was the acquisition interval for >70% of trajectories within the dataset.

### Classifying functional cell behaviors at single-cell resolution

With this curated dataset of cell trajectories (coordinates: *x, y, t*), we developed the CaMI framework to take the coordinates of cell trajectories, compute cell motility parameters using the anisotropic persistence walk (APRW) model^16^, perform *k*-means clustering analysis, and lastly identify and classify dynamic cell behaviors (**Figure 1A**). Using the 74,253 single cells, we computed eight motility parameters using the APRW model (**Figure 1B**, see **Methods**). Previously, we showed that the eight motility parameters generated using the APRW model were sufficient to capture the critical motility behaviors in a cohort of primary dermal fibroblasts^24^. Furthermore, using the 74,253 cell trajectories, we performed factor analysis to compare the communalities score of the eight APRW parameters with >30 other motility parameters computed based on Heteromotility^25^ (**Supplementary Figure 3A**). We found that the motility information captured by the eight APRW parameters (communality = 0.53) was greater than that of the >30 parameters from Heteromotility (communality = 0.39). Given this result, we chose to use the eight APRW parameters for further analysis. The APRW parameters include short and long time-scale mean squared displacements (MSD_10_, MSD_100_), persistence times in the primary and secondary axes of migration (P_p_, P_np_), total diffusivities and diffusivities in the primary and secondary axes of migration (D_tot_, D_p_, D_np_), and the anisotropic index that describes the spatial persistence (phi).

Using unsupervised *k*-means clustering and calculations of the inertia and silhouette values^26^ (**Supplemental Figure 3B**), we identified 17 distinct short-duration cell motility clusters (S1 to S17). To confirm this classification, we computed a coherence factor based on the vector distances between every cell and each cluster centroid **(Supplementary Figure 3C)**. Results showed the expected staircase plot, highlighting that cells within each cluster exhibited small distances (blue) to their assigned cluster centroids, and larger distances (white) to unassigned cluster centroids. Using two-dimensional t-stochastic neighbor embedding (tSNE), we plotted all 74,253 cells and painted each cell based on the designated motility cluster (n=17) (**Figure 1C**). We then assessed cluster groupings using unsupervised hierarchical clustering to describe gross similarities among cell motility clusters, resulting in four behavior classes: SG1, SG2, SG3, SG4 (**Figure 1D**). Each cluster and corresponding behavior class showed distinct motility patterns defined by the magnitude of the eight motility parameters (**Figure 1E-G, Supplementary Figure 3D**).

Overall SG1-4 can be summarized based on the following gross motility behaviors, respectively: SG1: low speed and high persistence; SG2: low speed and tortuous (low persistence); SG3: high speed and high persistence; and SG4: high speed and tortuous (low persistence). We also observed that the cell motility clusters belonging to SG1 and SG2 had lowed mean-squared displacements (MSDs) relative to motility clusters belonging to SG3 and SG4 (**Figure 1H**). Simultaneously, cell motility clusters in SG1 and SG3 had a higher persistence as denoted by a lower circularity score based on the probability density functions of their angular velocity magnitudes relative to SG2 and SG4 (**Figure 1I**). Circular profiled of the angular velocity magnitudes indicate that cells have a similar likelihood to move in multiple directions. To visually confirm that different clusters denoted distinct cell motility patterns, we plotted the trajectories of cells classified within each cluster, highlighting overt differences in cell behaviors per motility cluster (**Figure 1J**).

Together, this analysis demonstrates a multivariate approach to identify and classify distinct single-cell motility patterns across multiple datasets. All clustered trajectories and higher order parameters are reported in **Supplementary Tables 2 and 3**.

### Emergent short-duration motility patterns among distinct cell types and biological conditions

To determine how cell-types drive emergent cell behaviors, we quantified the abundance of cell trajectories across the 17 short-duration motility clusters for baseline and/or wild type conditions in various cell-types and performed unsupervised hierarchical clustering based on the magnitude of the correlation (**Figure 2A)**. Maintaining the same hierarchical clustering order by short-term trajectory, we repeated this quantification for the four short-duration behavior classes, SG1-SG4 and the eight APRW motility parameters. Results indicated that groups of cells and cell types present a range of heterogeneous motility behaviors, but certain cell types tended to cluster together. For example, immune cells formed one cluster (3D T-cell lymphoma line LY12, germinal center B-cells, 3D Jurkat T-cells and 3D T-cell lymphoma line Hut78), while fibroblast-type cells formed a separate cluster (2D and 3D fibrosarcoma, 2D human embryonic fibroblasts. Furthermore, even though groups of cells across conditions were enriched for one behavioral class over another (**Figure 2A center**), cells were generally biased towards unique combinations of short-term trajectories within each SG behavior class (**Figure 2A left**), suggesting heterogeneous phenotypes across cell groups. We also examined the same hierarchical clustering for a larger group of cell types with each group containing both baseline and molecular/environmental perturbations (**Supplementary Figure 4A**). Here, we see different clusters of cell types form, suggesting that these perturbations can shift baseline cell motility behaviors in such a way that cell types with distinct baseline conditions behave similarly in response to certain perturbations.

**Figure 2.**
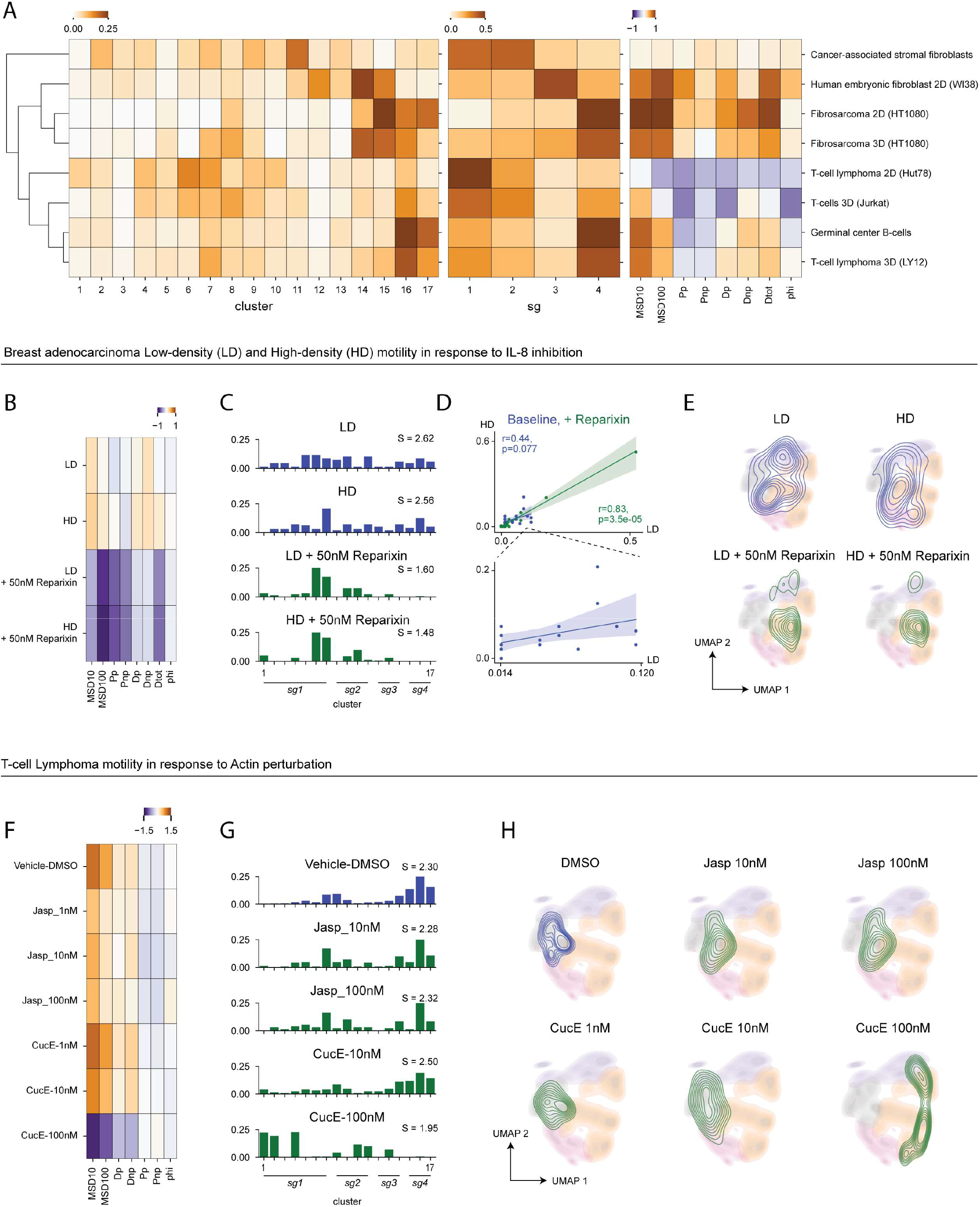
Application of CaMI workflow to short-duration datasets. **A**. Heatmaps indicating the fractional abundance of distinct cell types in our dataset across the seventeen short-duration motility clusters (left), four short-duration behavior classes (center), and the average magnitude of the eight motility parameters within each cell group (right). **B**. Heatmap of scaled higher order motility parameters for low-density (LD) and high-density (HD) MDA231 cells at baseline in response to 50nM Reparixin exposure. **C**. Barplots showing the fractional abundance of MDA231 cells within each of the seventeen short-duration motility clusters by condition. **D**. Linear regression plots indicating the correlation between LD and HD fractional abundance across the seventeen short-duration clusters at baseline and in response to Reparixin exposure. *R* and *p* indicate results from a Spearmen correlation test. **E**. Two-dimensional tSNE map with the four behavior classes shown as shadings and the contours of each MDA231 cell condition. **F**. Heatmap showing the magnitudes of the eight motility parameters for the LY12 T-cell lymphoma cells treated with Actin modulators (Jasplakinolide (Jasp.) and Cucurbitacin E (CucE)). **G**. Bar plots showing the fractional abundance of cells per short-duration motility clusters treated with Actin modulators. **H**. Contour maps showing the distributions and localizations of cells at baseline (DMSO) and treated with varying concentrations of Jaspand CucE.

Together, these results indicate that the changes in the overall magnitude of cell motility parameters are reflected by the redistribution of cells among motility clusters and behavior classes (**Figure 2A-C**). We note that while the CaMI platform harmonizes acquisition interval from different motility experiments, the current algorithm does not account for possible differences in culture conditions (i.e. cell density, type of well plate, etc.) between experiments that may impact motility behaviors. Therefore, similarities and differences between cell types reported here may still be confounded by these variables. As such, in subsequent sections we will directly apply the CaMI framework to profile cell motility patterns in specific test cases to highlight the biological insights that can be achieved relative to conventional cell motility analyses. Within each of these test cases, culture conditions between biological conditions were similar and enable us to make more direct conclusions about how certain perturbations impact motility behavior.

### Test case 1: Effect of cell density on emergent motility patterns of breast adenocarcinoma cells

To benchmark the ability of CaMI to yield biological insights, we first asked how this framework can elucidate emergent differences in cell motility in response to distinct environmental conditions, such as cell density. In a previous study, we showed that the MDA231 breast adenocarcinoma cells cultured at high density exhibited enhanced cell speed. Moreover, we showed that blocking IL-8 receptor signaling using the drug Reparixin can reverse this effect, suggesting that enhanced cell motility at high density conditions is due to paracrine IL8 signaling. Here, we were curious as to whether CaMI could further elucidate how Reparixin treatment impacts emergent single-cell motility *patterns*, that is collections of several bulk motility properties, beyond just displacement.

To test this hypothesis, we first extracted higher order motility parameters for control and 50nM Reparixin-treated cells cultured at low density (LD) and high density (HD) (**Figure 2B**). We found that on average, HD cells not only had greater displacement, but higher persistence time and diffusivity in the primary axis than their LD counterparts. When treated with 50nM Reparixin, the motility parameters shift such that both LD and HD cells have similarly low displacement, diffusivity, and persistence times. We then projected these cells onto the CaMI space (**Figure 2C**) and found that baseline LD and HD cells existed in distinct motility states of low- and high-speed. Within both density conditions, a subset of cells occupied the low displacement behavioral classes (SG1/2), while another subset occupied the high displacement behavioral classes (SG3/4). The baseline HD condition simply has a higher proportion of cells within the high-motility behavioral classes. This result shows that CaMI provides additional information about the heterogeneity of motility states within each cell condition. Instead of all single cells having HD in high-density conditions, CaMI enables us to see that different subsets of cells take on distinct behaviors. Treatment with Reparixin consistently skewed LD and HD cells towards high enrichment of SG1 motility patterns 6 and 7. Moreover, we quantified the Shannon entropy for each of these conditions and found that baseline conditions had higher entropy than those treated with Reparixin. This result indicates that Reparixin treatment also reduces the heterogeneity of motility states. Correlating the abundance of LD and HD cell types at baseline and in response to Reparixin treatment (**Figure 2D**) shows that these density conditions have stronger correlation in response to Reparixin exposure (Spearman *R* = 0.83, *p <0.001*) than at baseline (Spearman *R* = 0.44, *p = 0.07*). These results can also be visually demonstrated by projecting each cell type condition onto the CaMI short-duration tSNE space (**Figure 2E**).

Together, these results demonstrate how CaMI can confirm trends observed through bulk motility parameter calculation while providing additional information about the heterogeneity of motility states in response to environmental and molecular perturbations.

### Test case 2: Effect of cytoskeletal modulators on T-lymphoma motility in 3D microenvironments

Next, we were curious whether CaMI can quantitatively demonstrate how perturbing cellular mechanisms that directly drive cell movement, such as the cytoskeletal network, shift cell motility behaviors. To explore this question, we applied CaMI to test the effect of actin and microtubule modulation in T-lymphoma cells (LY12) within 3D collagen-I microenvironments. To test the effects of actin and microtubule modulation on motility behavior, we exposed cells to escalating concentrations of four drugs: Jasplakinolide (Jasp), an actin polymerizer, Curcubitacin E (CucE), an actin depolymerizer (**Figure 2F-G)**, Taxol, a microtubule stabilizer, and Vincristine (Vinc), a microtubule depolymerizer (**Supplementary Figure 4B-D**).First, we quantified the bulk motility parameters for each cell type condition (**Figure 2F, Supplementary Figure 4B**). At baseline LY12 cells have high displacement and low persistence times. We observed that increasing Jasp concentration slightly decreaed cell motility, as noted by the slight drops in motility parameters like Dtot and MSD10 (**Figure 2F**). However, increasing concentrations of CucE appeared to have a much more significant decrease in cell displacement, as indicated by the stronger drops in the values of these motility parameters (**Figure 2F**). Using the CaMI workflow, we observed that baseline LY12 cells occupied SG1 cluster 6/7 or behavior class SG4 clusters 14-17 with a stronger enrichment towards SG4. Treatment with increasing concentrations of Jasp appeared to slightly increase enrichment in SG1 clusters, while decreasing that in SG4. LY12 cells treated with increasing concentrations of CucE, on the other hand, had a much stronger response with cells being depleted from the SG4 behavior and enriched in either SG1 or SG2. Here, CaMI provides additional information that bulk motility parameter analysis by showing that the drop in displacement parameters in response to 100nM CucE treatment was reflected by a divergence towards two distinct cell motility states. The Shannon entropy for the cells treated with 100nM CucE was also much lower than at baseline or in response to Jasp, suggesting that CucE reduces the heterogeneity in cell motility as well. These trends are also visually reflecte din the short-duration motility tSNE space (**Figure 2H**) where we observe the shift of cells treated with 100nM CucE from the left to right of the low-dimensional space.

Interestingly, exposures to microtubule stabilizer Taxol led to a similar response as exposure to Jasp, wherein cells experience a slight decrease in displacement and enrichment in SG1 motility patterns (**Supplementary Figure 4B-C**). Conversely, treatment with Vincristine led to a strong increase in cell displacement from baseline and greater enrichment in SG4 motility patterns (**Supplementary Figure 4B-C**). These patterns were visually observed by the contour of cells treated with Taxol slightly expanding in the low-dimensional motility space, while contour of cells treated with Vincristine progressively shrinks within SG4 (**Supplementary Figure 4D**).

Together, these results indicate how the CaMI framework can quantitatively demonstrate cellular heterogeneity that are not directly apparent using conventional cell motility analyses. Particularly, CaMI can elucidate instances where change in a bulk motility parameter translates to enrichment of two distinct motility states. Treatment with 100nM CucE decreased motility parameters such as MSD10 and MSD100, but this decrease translated to enrichment of two distinct behavioral classes—SG1 and SG2. This result means that the drug causes cells to take on both low displacement, high persistence and low displacement, low persistence motility behaviors.

### Using CaMI to predict cell motility clusters in previously unclassified cells

In addition to using CaMI to decipher how molecular and environmental perturbations drive fractional redistributions across motility states, we wondered whether we could project new data into our dataset to quantify behaviors for previously unclassified cells. To perform this experiment, we trained a linear support vector machine learning model (90:10 train-test split) to predict short-duration motility pattern (S1-S17) based on the underlying eight APRW parameters of individual trajectories. Our trained linear SVM had an average training accuracy of 98.7% across all clusters (**Supplementary Figure 5A**). To validate the model, we simulated 75,000 new trajectories using a model developed by Wu et al^16^. With these new trajectories, we computed the 8 motility parameters and performed the classification using the trained 90:10 model, which yielded an average validation accuracy of 96.7% across all clusters (**Supplementary Figure 5B**). Here, we validated the true motility cluster of each simulated cell by performing an iterative analysis to re-classify all the cells adding one new simulated/unclassified cell each time (*i.e*., re-training on 74,253 cells + 1 simulated cell; 75,000 times). Having demonstrated a high accuracy of the trained model, we tested the model using cell trajectories of primary monocytes moving within 1mg/ml Collagen-I gels.

### Test case 3: Fractional redistributions among cell motility clusters define monocytes motility

Having trained a model to now project new cell measurements onto the CaMI motility space, we were curious as to how CaMI can quantitatively assess cell motility properties in primary human cells. To explore this question, we specifically focused on primary human monocytes because these cells are known to rely on motility to orchestrate immune function in response to local microenvironmental cues and signals. Moreover, prior studies have shown the aging and cancer can disrupt monocyte ability to perform these critical functions, suggesting that having a tool to assess the ability of primary monocytes to navigate cellular environments can help assess patient health and disease potential. With the trained linear SVM model, we quantified the distributions of short-duration motility patterns for primary monocytes that were not included in the initial CaMI workflow and dataset. Briefly, we collected peripheral blood mononuclear cells (PBMCs) from three healthy donors (D1, D2, D3) and isolated monocyte populations based on CD14+ expression. We then embedded monocytes from each donor into 1mg/ml Collagen-I gels, with eight technical replicates (*i.e*., wells in a 96 well plate) per donor. In half of the wells per donor (4 wells each), we exposed monocytes to conditioned media from breast cancer cells (MDA-MB 231) for approximately 12 hours prior to imaging. Using 3D time-lapse microscopy, we imaged and analyzed the monocytes moving in three-dimensional collagen gels (2mg/ml) for 2.5 hours at 5-minute intervals then applied the aforementioned linear SVM to classify the motility pattern of each cell.

We observed that each donor exhibited unique baseline profiles of short-duration motility patterns (**Figure 3A**). Donor 3 exhibited high baseline abundances of cells in the SG4 behavior class (specifically in motility clusters S16, S17), while donors 1 and 2 exhibited higher abundances of cells in SG2 behavior class. Furthermore, when monocytes were exposes to conditioned media (+CM), donors 1 and 2 exhibited similar shifts towards enrichments in behavior class SG4, specifically motility clusters S14, S15, S16, S17, corresponding to high displacement and low persistence motility behaviors (**Figure 3A**). These shifts in cell motility patterns after CM exposure were also associated with decreased cellular heterogeneity as determined by the Shannon entropy (**Figure 3B**) and a higher correlation seen among cells exposed to CM relative to their baselines (**Figure 3C**). Interestingly, donor 3 did not exhibit drastic changes in the distribution among cell motility clusters after CM exposure, with no significant observed in the baseline and CM treated cells based on the total diffusivity (*i.e*., effective speed) (**Figure 3D**). Together, the results confirm that monocytes exposed to CM-associated signals increase their movements, with the lack of response in donor 3 suggesting a resistance or insensitivity to CM-associated signals. While the magnitude of the response can be determined using conventional cell motility analysis, the addition of CaMI analysis provided a direct visualization of the responses and demonstrates that the ensemble response itself (increased cell motility) is due to a fractional redistribution among cell motility clusters.

**Figure 3.**
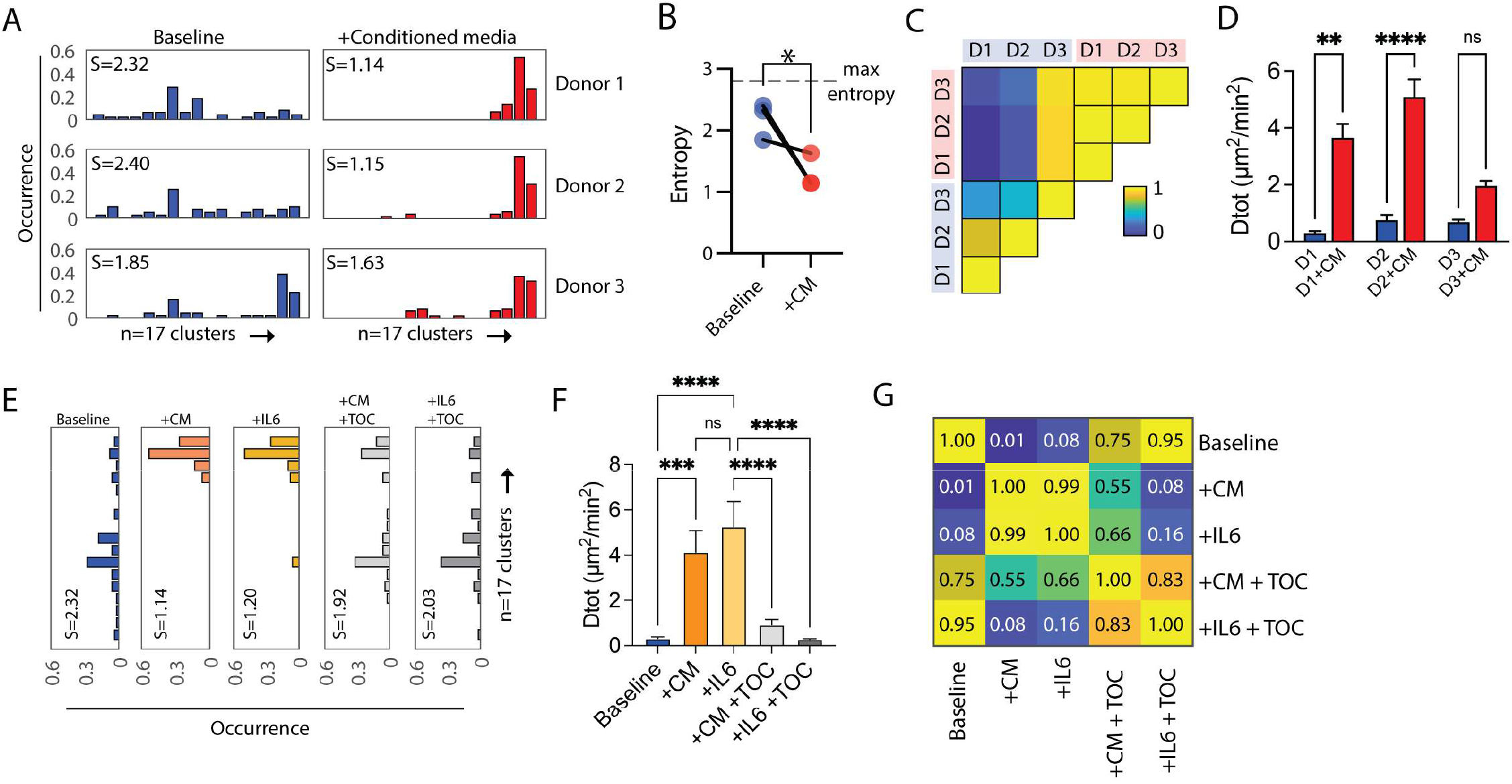
Quantification of cell motility patterns in primary monocytes. **A**. Bar plots showing the abundance of cells among the 17 cell motility clusters for three donors at baseline and after exposure to conditioned media. **B**. Scatter plot showing the decrease in cellular heterogeneity after CM-exposure. **C**. Correlation map of cell motility clusters among donor samples. **D**. Bar plots showing the magnitude of the total diffusivities at baseline and after CM-exposure. **E**. Bar plots showing the distributions of cell motility clusters across testing conditions. **F**. Bar plot showing the total diffusivities across testing conditions. **G**. Correlation map indicating similarities in the cell motility clusters among testing conditions.

To decipher these responses further, we asked whether specific molecules within the conditioned media were capable of driving similar shifts in the motility behavior. Since cells from donor 1 exhibited drastic shifts in response to CM exposure, we selected that sample to performed additional experiments. Recent studies on cancer cell conditioned media indicates that interleukin-6 (IL6) is highly expressed^27^. Furthermore, other studies have shown that IL6 exposure increases cell migration in monocytes^28^. To test the effects of IL6 exposure on monocyte motility patterns, we exposed cells from donor 1 to either CM or 500ng/ml of recombinant human IL6 and profiled cell motility patterns. Similar described above, cells exposed to both CM and IL6 exhibited a characteristic shift in the abundance of cells among cell motility clusters to enrich for cells in behavior class SG4 (specifically S14, S15, S16, and S17), with a corresponding decrease in cellular heterogeneity and higher total diffusivities/effective speeds (**Figure 3E-F**). Lastly, to test whether IL6 was directly modulating the phenotype, we co-treated cells with either conditioned media or IL6 and 100μl IL6-receptor blocking antibody Tocilizumab. Results indicated that monocytes treated with Tocilizumab regained a baseline-like profile denoted by the shifts in the distributions among cell motility clusters (**Figure 3E**), a decrease in total diffusivity, and an increase in the magnitude of the cellular heterogeneity to baseline levels (**Figure 3F**). Furthermore, correlation analysis on the distributions of cells across cell motility clusters among conditions showed that cells exposed to CM and IL6 were highly correlated, with baseline monocytes showing strong correlation with cells co-treated with CM or IL6 and Tocilizumab (**Figure 3G**).

Altogether, we show that using the CaMI framework, we can directly quantify dynamic cell behaviors, and cellular heterogeneities to profile emergent responses of cells using cell motility. Furthermore, results indicate that all the biological insights gained from conventional motility analyses are conserved, with new insights gained regarding heterogeneity, fractional redistributions, and the direct visualization of the cell motility trends at baseline and after perturbations.

### Quantification of dynamic cell behaviors over long durations

In many of the conditions used, more than eight hours of continuous cell tracking data was available. We wondered what long-duration motility patterns existed within this motility database, and how these long durations motility patterns related to the short-duration cell motility patterns (2.5 hours). We rationalized that this could be informative, especially since current approaches to analyze cells motility relies on the assumption of time invariance. To evaluate these questions, we compiled cells with eight hours of motility data at 5-minute intervals and computed the same eight motility parameters described in the short-duration analyses. With this data (n=24,753 cells) (**Figure 4A**), we performed *k*-means clustering analysis to identify 25 clusters (**Figure 4B**). We verified cluster number using the inertia and silhouette values and coherence staircase plot (**Supplementary Figure 6A-B**). These 25 motility clusters (C1 to C25) also separated into four behavior classes (CG1, CG2, CG3, CG4) based on unsupervised hierarchical clustering using correlation distances (**Figure 4C**). Each long duration motility pattern and behavior class displayed distinct patterns in the magnitudes of the eight motility parameters (**Figure 4D-E, Supplementary Figure 6C-E**). Plotting the trajectories of cells within each of the long-duration motility clusters revealed not only distinct patterns of movements (**Figure 4F**) but also a polarization of highly motile cells defined by MSD100 in motility clusters belonging to behavior classes CG3 and CG4 (**Supplementary Figure 6C**) and higher persistence for cells in behavior classes CG2 and CG3 (**Supplementary Figure 6D**).

**Figure 4.**
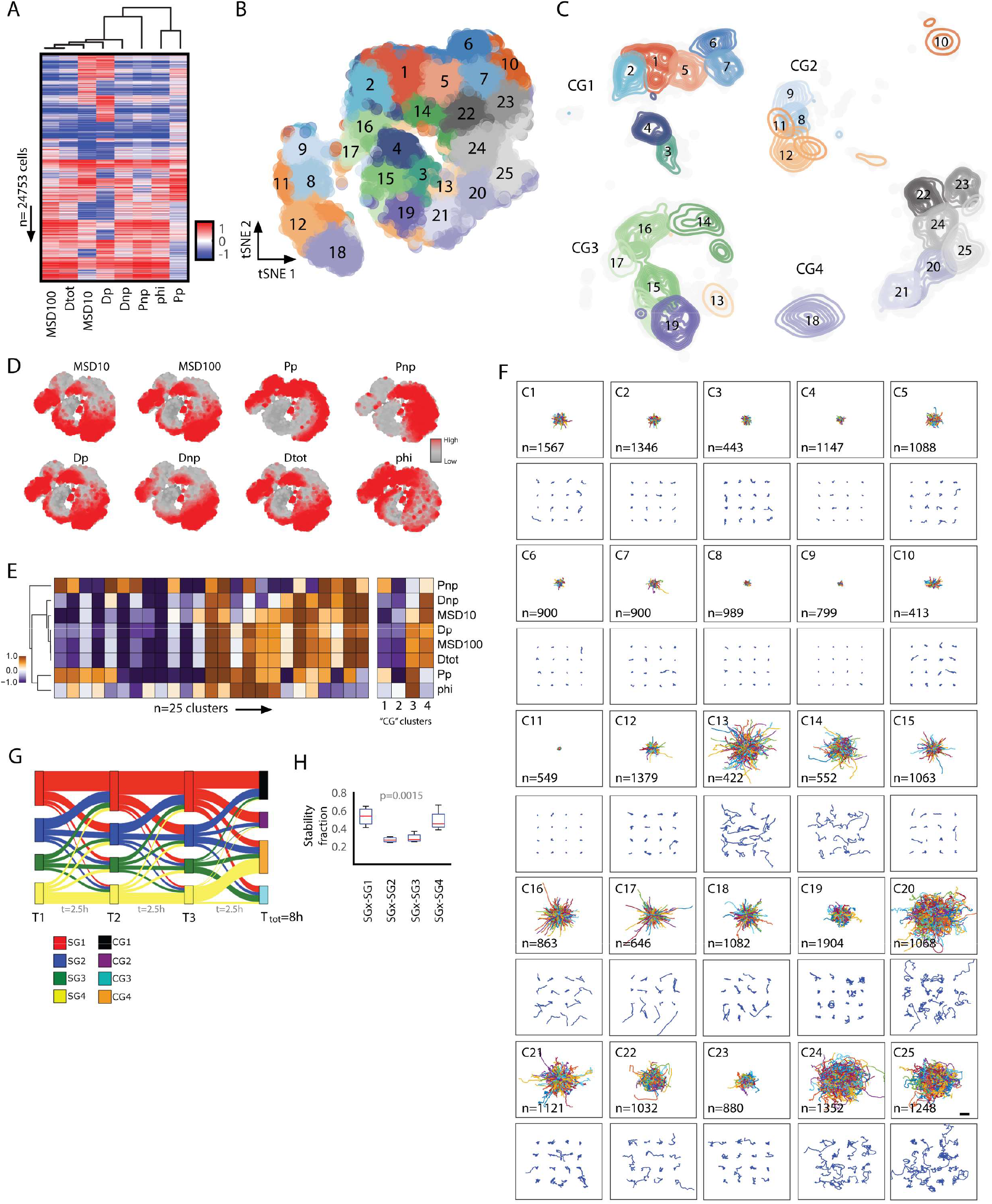
Identifying and classifying cell motility patterns across long durations. **A**. Heatmap showing eight motility parameters per cell (n=24,753 cells). Each row represents a single cell, and each column represents a motility parameter. **B-C**. Two-dimensional t-SNE scatter plot where each dot is a cell and colors indicate the 25 identified long-duration cell motility clusters (B) and contour plots further separated into four long-duration behavior classes (CG1-4) that show the distributions and localization of cells per motility cluster (C). **D**. Two-dimensional t-SNE plot showing the magnitude of the eight motility parameters per cluster region. **E**. Heat map showing the magnitude of the eight cell motility parameters for the twenty-five long-duration cell motility clusters (left) and within each of the four long-duration behavior classes (right). **F**. Visualization of cell trajectories for cells classified within each cluster, the top panel shows the origin-centered trajectories for all cells within a cluster, and the bottom panel shows 16 randomly selected cells per cluster. Scalebar denotes 100μm. **G**. Sankey plot showing the transitions based on the fractional abundance of cells within each short-duration behavior class for three-time segments (2.5 hours each) and how they ultimately connect to the four long-duration behavior classes. **H**. Box-and-whisker plots of the stability fraction of cells within each short-duration behavior class.

We next evaluated the ability to predict the motility pattern of unclassified cells. Using the same approach described for the short-duration motility clusters. Similarly described above, we trained a new linear SVM model on the 8-hour cell tracking data, tested the prediction in a 90:10 split dataset, and newly simulated trajectories for 25,000 single cells. Results showed a 97.2% accuracy for the 90:10 split dataset (**Supplementary Figure 7A**) and a 95.6% accuracy for the simulated trajectories (**Supplementary Figure 7B**). With this tool, we can also identify and classify distinct cell long-duration motility patterns for cells not used to train the CaMI platform (clustered long duration trajectories and higher order parameters reported in **Supplementary Table 4-6**).

### Connecting motility patterns across short and long durations

In most cell motility analyses, the inherent assumption of time-invariant movement or time independence limits the ability to infer temporal stability of cell behaviors. To infer temporal patterns, we coupled short and long-duration motility clusters within the same cells to determine temporal transitions between short-term motility behaviors, thereby inferring overall long-term temporal patterns of motility behaviors. For a subset of cells having 8 hours of movement, we broke each 8-hour trajectory into three non-overlapping 2.5-hour segments. We determined motility clusters (S1-17 and C1-25) and complimentary behavior classes (SG1-SG4 and CG1-CG4) for all short-duration segments as well as the long-duration trajectories (**Supplementary Table 2, Supplementary Table 5**). With this data, we then asked whether a cell classified in a particular motility cluster or behavior class for the first 2.5 hours stayed within the same short-term behavioral class for subsequent 2.5-hour time segments. Overall, we observed that for the ∼25,000 cells assessed, cells classified within classes SG1 and SG4 tended to stay within the same behavior class for the three consecutive time segments (**Figure 4G**), with higher temporal stabilities across the time segments (**Figure 4H**). Conversely, SG2 and SG3 tended to shift behavior classes over time and had relatively lower temporal stability.

Taken together, we describe the ability to quantify the temporal dynamics of the motility patterns at short and long durations. This quantification of temporal dynamics is critical, as it provides a robust approach to evaluate temporal cell motility patterns without the direct limitations of the time invariance assumption. By separating cell trajectories into shorter time segments, where the time-invariance assumption is valid, we can decipher spatio-temporal patterns that accurately describe single-cell behaviors.

### Using CaMI to evaluate functional heterogeneity across conditions

Given the ability to profile dynamic behaviors of single cells using CaMI, we further tested the ability to derive key biological insights that were not directly apparent using conventional motility analyses based on average motility parameters alone. Using a subset of data from long-duration movies (*i.e*., 8 hours), we performed hierarchical clustering based on cross-correlation analysis across the 409 conditions. This analysis yielded four groups, as shown in the cross-correlation heatmap (**Figure 5A**) and two-dimensional tSNE plots (**Figure 5B**). As a biological test case, we took data from LY12 cells (T-cell lymphoma cells) moving within 3D collagen-1 gels across five increasing collagen concentrations. Plotting the trajectories of cells within each condition, we qualitatively observed a bi-phasic response in the overall magnitude of the cell displacements, with the lowest displacements observed at 3mg/ml collagen (**Figure 5C**). Plotting this condition in the 2D t-SNE space, we quantitatively observed a bi-phasic or switching effect (cells in 4mg/ml are more similar to cells in 1mg/ml than to cells in 3mg/ml (**Figure 5D**). This switching phenotype with changing collagen concentrations is likely due to local changes in molecular crowding effects and effects on collagen alignment and porosity^29,30^. Furthermore, plotting the abundance of cells within the 25 clusters for each condition, we also observed a bi-phasic response in the heterogeneity (defined based on the Shannon entropy), with a minimization at 3mg/ml (**Figure 5E**).

**Figure 5.**
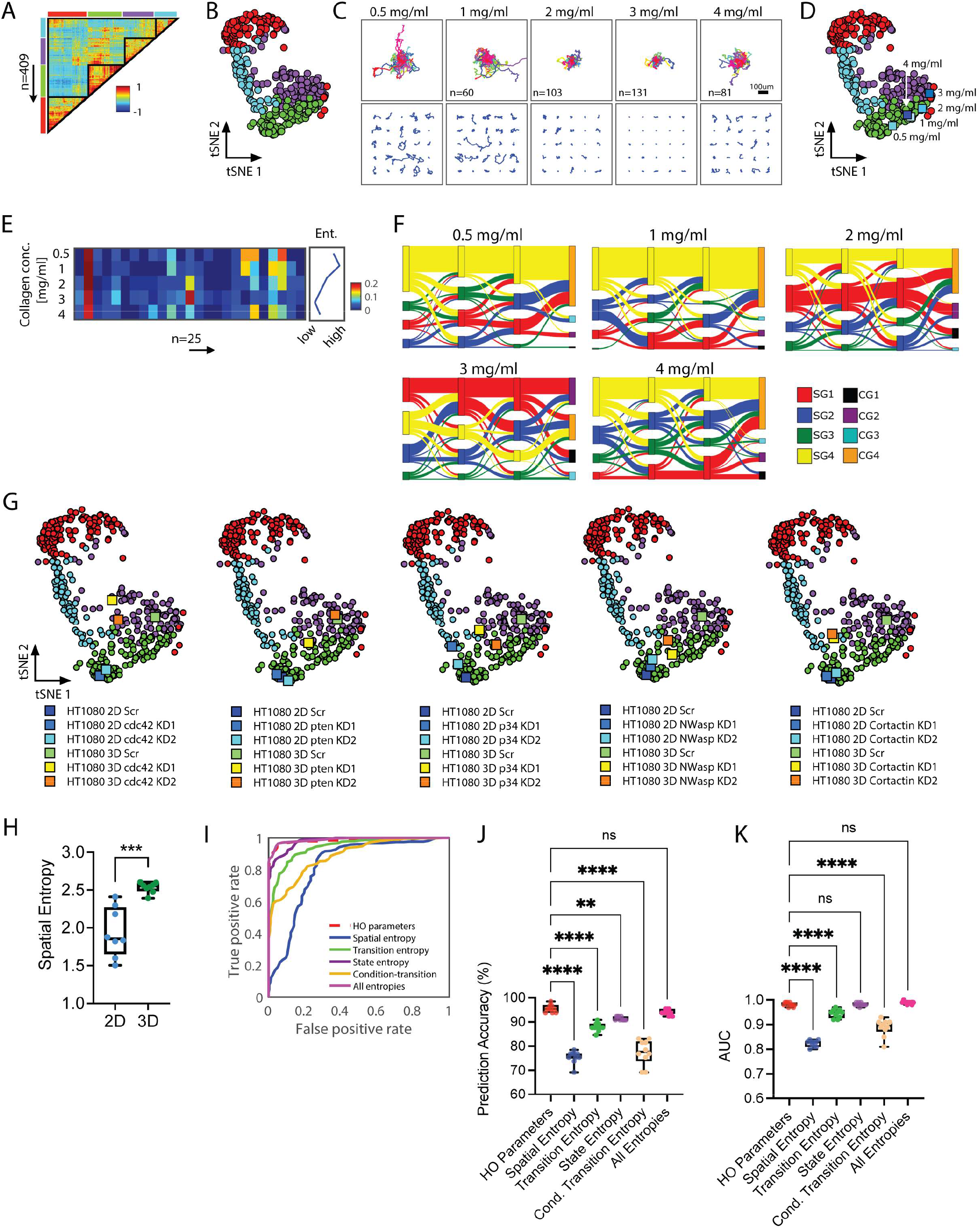
Quantifying functional heterogeneity and biological insights using CaMI. **A**. Cross-correlation heatmap highlighting the grouping of the 409 biological conditions based on similarities in overall cell behaviors. **B**. Two-dimensional t-SNE plot showing the 409 conditions painted based on the identified groupings. Each dot denotes a single biological condition (n=409). **C**. Single-cell trajectories of LY12 cells as a function of increasing collagen concentrations. The top panel shows origin-centered trajectories for all cells analyzed, and the bottom panel shows 25 randomly selected cells per condition. **D**. Two-dimensional t-SNE plot shows a bi-phasic/switching effect with increasing collagen concentration. LY12 cells moving in 4 mg/ml gels are more similar to LY12 cells moving in 1mg/ml gels relative to cells moving in 3mg/ml collagen-I gels. **E**. Heatmap showing the abundance of cells within each cluster per collagen concentration. The line plot on the right shows a bi-phasic change in the Shannon entropy, denoting the lowest cellular heterogeneity for cells moving in 3mg/ml collagen-I gels. **F**. Sankey plots show the temporal transitions of cells across short-duration classes as a function of increasing concentration. Sankey plots also indicate a switch in temporal kinetics and temporal heterogeneity. **G**. Two-dimensional t-SNE plot showing the location of matched 2D and 3D conditions of ht1080 cells. **H**. Box plot showing the Shannon entropy of cell moving in 2D microenvironments are significantly lower than for cells moving in 3D microenvironments. **I**. ROC curve showing prediction accuracy, denoted by the true and false positive rates, for higher order motility parameters and different spatio-temporal entropies. **J-K**. Box- and-whisker plots show the prediction accuracy (J) and area under the curve (K) for higher-order motility parameters and various spatio-temporal entropies. Data indicates that combining entropies yields the same prediction accuracy as using the higher-order parameters.

Given these striking spatial associations with collagen concentration, we asked whether complementary temporal associations were based on short and long durations. To address this, we stacked the three 2.5-hour time segments, paying attention to the fraction of cells within each short-duration behavior class (SG1 to SG4) and the frequency at which cells switched classes with increasing concentrations. From this analysis, we also observed a switching phenotype based on temporal associations, with the majority of cells staying in SG4 (yellow) across the three segments for 0.5mg/ml, 1mg/ml, and 4mg/ml, an increase towards SG1 for 2mg/ml, and a shift towards SG1 as the dominant behavior class for 3mg/ml (**Figure 5F**). Furthermore, while this phenotype switch is observed spatially at 3mg/ml, from the temporal perspective, this shift is already apparent at 2mg/ml and is enhanced at 3mg/ml.

Altogether, CaMI enables the quantification of real spatial-temporal dynamics and heterogeneities that define bi-phasic and phenotype-switching responses to increasing collagen concentration.

### Predicting dimensionality of cellular conditions based on spatial and temporal heterogeneities

Given the ability to dynamically map spatio-temporal cell behaviors, we wondered whether we could predict the dimensionality of cells moving in 2D and 3D microenvironments. Recent studies have demonstrated that cell movements in 2D do not necessarily recapitulate 3D movements^19,31,32^, which is especially applicable in studies involving the characterization of metastasis and cancer cell invasion. Furthermore, it is unclear why there is such a disconnect in motility behaviors in 2D and 3D microenvironments, where 2D cancer cell migration is not predictive of cancer cell invasion. We evaluated cell motility patterns in 60 matched 2D and 3D conditions to elucidate what defines this difference in 2D vs. 3D. These conditions were comprised of ht1080 cells under multiple genetic and drug-induced perturbations, including knockdowns using different shRNAs and concentration-dependent drug treatments. This wide range of perturbations allows direct interrogation of the universe of movement patterns exhibited by cells moving in 2D and 3D microenvironments. Mapping the 2D and 3D conditions in t-SNE space, we observed differential localization across regions (**Figure 5G**), indicating inherent differences in the motility patterns of the cells, with cells in 3D microenvironments exhibiting higher heterogeneities (**Figure 5H**). This differential localization within the 2D t-SNE space was observed in knockdown conditions and as a function of drug concentrations in ht1080 cells and other cell types (**Supplementary Figure 8**). Although we did not observe dimension-exclusive motility clusters (meaning that both 2D and 3D cellular conditions had cells classified in each of the 25 long-duration motility clusters), the fraction of cells within each cluster varied greatly between 2D and 3D (**Supplemental Table 2-3, Supplementary Table 4-5**).

Prompted by these results, we asked whether we could predict the dimensionality of moving cells based on the magnitude of the heterogeneity across the cell populations since cells in 3D tended to exhibit higher phenotypic heterogeneities relative to matched 2D conditions (**Figure 5H**). To comprehensively define the heterogeneity, we further defined four types of entropies that captured cells’ spatial and temporal nature (see **Methods**). These different types of entropy were (a) spatial entropy, which is computed based on the fractional abundance of cells within each short or long-duration motility cluster (n=17 or n=25), (b) state entropy, computed based on the fractional abundance of cells within short and long duration behavior classes (SG1-SG4, CG1-CG4), (c) transition entropy, which is computed based on the probability of transitioning from one short duration behavior class to the another along subsequent time segments, and (d) conditional-transition entropy, estimates the entropy based on the conditional probabilities that define the likelihood of transitioning to a specific short duration behavior class. We then trained a model to predict the dimensionality of cells using a linear SVM for each entropy and the combination of entropies (see **Methods**). Comparing the prediction accuracies using either the eight motility parameters or the entropies, we found that the combination of entropies was able to predict the dimensionality in more than >95% of the cases, which was comparable to using the eight motility parameters to predict the dimensionality (**Figure 5I-K**).

Taken altogether, we show the ability of CaMI to predict patterns of cell movements and derive biological insights across cellular conditions, such as how the dimensionality of the microenvironment influences dynamic behaviors. Furthermore, understanding phenotypic heterogeneity (both spatial and temporal) is critical to interpreting single-cell and population-level cell behaviors.

## DISCUSSION

Cell movements play a critical role in the physiology of organisms, both in homeostasis and disease^33^. CaMI is a computational framework that takes advantage of the single-cell nature and spatio-temporal dynamics of cell motility data to quantitatively extract distinct motility patterns across multiple conditions. Conventionally, cell motility is quantified using small numbers of cells and reported as bulk averages of cell populations. Here, we have demonstrated the utility of CaMI to quantify cell motility patterns, determine spatial and temporal heterogeneities, define dynamic phenotypic transitions, and predict the dimensionality of cellular conditions. In addition, CaMI provides a direct visualization scheme of single-cell behaviors to enable direct interpretation of motility data while retaining all the biological information gained from conventional bulk analyses. Furthermore, CaMI can classify dynamic cell behaviors from imaging data extracted from either *in vitro* and *in vivo* settings or response to genetic or chemical perturbation, *e.g*., drugs and shRNAs.

While CaMI provides a robust solution to decipher emergent patterns of single-cell behaviors, there are certain contexts where it may be limited. A fundamental limitation of CaMI is that dominant cell motility patterns would be identified as clusters since cell motility clusters are derived based on multivariate statistical approaches of combined motility datasets (*i.e*., spanning multiple cell types, dimensionalities, and perturbations). As a result, rare, cell-type-dependent, or perturbation-dependent cell motility patterns may not be identified as distinct but will be classified within larger, dominant clusters. While this is the case in most big-data classifications, CaMI can provide reliable and insightful information, as demonstrated by the high cluster coherence for cell-type-dependent or perturbation-dependent conditions by retraining the motility clusters using an appropriate dataset. In conditions where cell type or condition-specificity is critical. One way around this limitation is to train the classification model based on a specific cell type or condition. For instance, we have done this for dermal fibroblasts that resulted in eight total motility clusters ^7^. However, the size of the dataset (*i.e*., the number of cells) must be large enough to be robust and reproducible. Lastly, CaMI is designed as a comparative tool, where cellular conditions are differentiated based on the fractional redistribution among defined cell patterns. Multiple comparative conditions are needed to derive deep biological insights and understand how a given perturbation influences the resultant cell behaviors based on the abundance of cells within each motility cluster or behavior class.

Additionally, the test cases presented in this study came largely from in vitro experiments of cells moving on two-dimensional substrates or within three-dimensional hydrogels. While we only looked at (x, y, t) coordinates, the framework can be expanded to look at true 3D movements (x, y, z, t) in addition to 2D movements of cells in a 3D hydrogel. Lastly, the CaMI framework can be used on cells moving in vivo based on movies from intra-vital imaging or cells moving in ex vivo organotypic cultures (viable sections of thick tissues that can be mounted onto a microscope). In short, as long as the coordinates (x, y, t) or (x, y, z, t) can be generated, the CaMI workflow can be applied and tested, the key consideration is the dataset of cell trajectories that are used to identify the cell motility clusters.

Altogether, we present a robust framework to identify and classify dynamic single-cell behaviors across multiple conditions. We anticipate that increased implementation of modern single-cell approaches and data science analyses will lead to a better understanding of dynamic cellular behaviors and heterogeneity. While CaMI provides a robust approach to quantify functional cell behaviors, additional work is needed to better understand phenotypic transitions and mechanistic associations of cell behaviors to their underlying molecular states. Recently, studies have started to multiplex dynamic properties of cells with underlying genetic and epigenetic states of cells^34–36^. However, more studies are needed to comprehensively assess and establish deterministic links between cell motility patterns and behaviors, cell state transitions, cell fates, and underlying molecular modifications (*e.g*., cells with knockdowns in particular genes). Ultimately, multiplexed *in situ* quantification of molecular and phenotypic behaviors quantified simultaneously could enable precision health approaches where dynamic cell behaviors connect to disease states. This link will be critical in light of heterogeneous cellular behaviors within cell populations and the associations with clinical outcomes and treatment responses^37,38^.

## MATERIALS and METHODS

### Data Curation

We compiled previously published data and newly acquired data to generate an *in vitro* single-cell motility database for the following analyses ^7,17–22^. All motility data were derived from time-lapse movies collected using brightfield or phase microscopy. Single-cell trajectories were extracted using commercial cell tracking software (Metamorph). Cell motility data were compiled from several different sources, with movies ranging in total duration from 2.5 to 16 hours and acquisition intervals from 2 to 10 minutes. All trajectories were collected in 2 dimensions (x, y); the database contains experiments of cells moving both within 2D substrates or 3D scaffolds, but the analyses here only consider movement in two dimensions for both cases. The entire database comprises of 74,253 single cells across 409 unique biological conditions. **Supplementary Table 1** contains the total duration, interval time, pixel size for calibration, and experimental descriptions for each biological condition. To render the individual single-cell trajectories comparable, we applied statistical corrections (described below as pseudo-Monte Carlo) to generate single-cell trajectories with consistent acquisition intervals of 5 minutes. From these videos, we generated short-duration trajectories of 2.5 hrs and long-duration trajectories of 8-hour duration. To generate these trajectories, longer videos were cut into shorter sections meaning that multiple trajectories in our training data could pertain to the same cell. We harmonized 74,253 short-duration, 2.5-hour trajectories (**Supplementary Table 2**) and 24,753 long-duration, 8-hour trajectories (**Supplementary Table 4**).

### Assessing geometric concordance among varied acquisition time intervals

To understand how acquisition time interval impacts trajectory shape, we selected ht1080 fibrosarcoma cells moving within 3D Collagen-1 gels because this dataset was initially acquired at the highest temporal resolution of 2 minutes. We *in-silico* generated degraded time-resolution trajectories for each original trajectory with acquisition times ranging from 2.5 to 10 minutes in 0.5-minute increments. We measured the concordance between trajectories with varied acquisition times. The concordance (**Supplementary Figure 1A**) was quantified as the summed triangular area formed between points acquired at consecutive timesteps.

### Pseudo-Monte-Carlo technique to predict x,y locations of cells at a given time point

First, we estimate the required (or missing) X-Y location from linear interpolation ( *x*_int *erp*,*i*_, *y*_int *erp*,*i*_ ) . Then, we employ a small perturbation to the interpolated point according to the equations below:

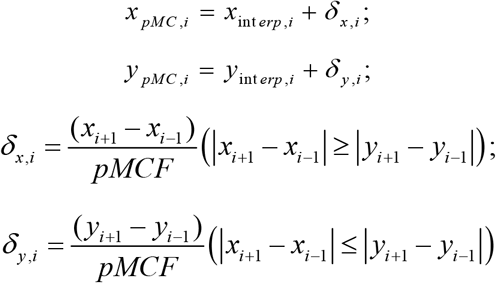

The perturbation is either applied to the dimension that experiences more displacement in a given interval. For example if between two time steps the cell moves more horizontally than vertically, the perturbation is applied in the X dimenion and no perturbation is applied in the Y dimension. A perturbation is applied in both dimensions when the cell moves perfectly horizontally. We tested *pMC factor* values between 2 and 2.03 (**Figure S1C)** and statistically compared resultant higher order parameters using a Kruskal-Wallis test followed with the Dunn test for multiple comparisons testing. We selected a *pMCF* value of 2.01 because this yielded no statistically significant difference from the original trajectory for all higher order parameters. A *pMC* factor of 2.0 was also sufficient but there were certain situations where *pMC* factor 2.01 performed better.

### Calculation of Higher Order Parameters using anisotropic persistence random walk (APRW)

To quantify the spatio-temporal motility patterns, we analyzed bulk cell movements based on trajectories using the Anisotropic Persistent Random Walk model (APRW)^19,23^. From this analysis, we generated the parameters that describe the movements of the cells, namely, MSD10, MSD100, Dtot, Dp, Dnp, Pp, Pnp, and ϕ. We calculated MSD10 and MSD100 since the denoted regimes are shorter and longer than the average persistence time across all conditions. We used the following equations to estimate the motility parameters.

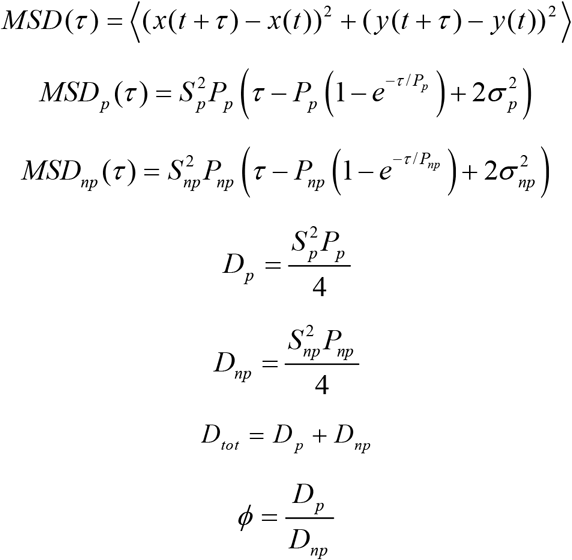

where *S* is the cell speed, *P* is the persistence time, 2σ^2^ is the noise (error) in the position of the cell, *τ* = *n* Δt and *n* = 1, 2, … N_min_−1, Δt is the size of the time step.

### Parameters and descriptions

The eight motility parameters used for all analyses include the magnitudes of cellular displacements and speeds (MSD10, MSD100), the total diffusivity and diffusivities along primary and secondary axes of migration (Dtot, Dp, and Dnp), the persistence times along the primary and secondary axes of migration (Pp, Pnp), and the spatial persistence/anisotropy (φ). Time lags of 10 and 100 (for the MSDs) were selected based on the average persistence times across datasets. Ten minutes were on the order of average persistence time, and 100 minutes was greater than the average persistence time.

### Normalized Mean Squared Displacement (MSD) Calculation

We normalized the Mean Squared Displacement (MSD) per time point to their corresponding time interval. For the non-homogenized trajectories, we used the following equation:

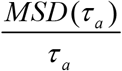

Where *τ*_*a*_ is the time interval of the actual non-homogenized trajectory. We used pseudo-Monte-Carlo to extract the normalized MSDs at varying time intervals as follows:

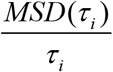

*τ*_*i*_ is the time interval used (*τ*_*i*_*=*2, 2.5, 3, 3.5, … 10 min).

### Clustering analysis, evaluating cluster stability and cluster coherence

We performed k-means clustering on the correlation distances to determine the emergent motility clusters among short-duration and long-duration trajectories. To determine behavioral classes of trajectories, we performed unsupervised hierarchical clustering on the centroids of k-means identified clusters, using the average linkage method and correlation distance metric. To analyze cluster stability, we used two factors: Inertia and Silhouette Values. Inertia values were based on the elbow method to identify the number of clusters when minimal additional data is gained with increased clusters^26^. We calculated silhouette values in MATLAB’s silhouette function. To compute cluster coherence, we computed the Euclidean distances between the z-score, log normalized motility parameters for each cell, and z-score, log-normalized motility parameter values of the centroid of each cluster (obtained through k-means clustering).

### Simulation of Trajectories

To simulate trajectories, we used APRW-based simulation of trajectories, as detailed in Wu et al. ^23^. To test classification accuracies across short- and long-term durations, we simulated 75,000 and 25,000 unique cell trajectories, respectively.

### Support Vector Machine (SVM) based data classification and model prediction

We used MATLAB’s built-in SVM (ClassificationsSVM) function to train on datasets and predict from the trained model. The trained model was a linear SVM using a hinge loss function with 90:10 train-test split. To get the cluster of a single cell, first, we calculate the higher-order parameters of the cell. Then we performed z-score and log normalization to receive the input for a single cell to the SVM-based classification.

### Centroid based prediction

To validate SVM-based prediction, we used centroid based classification system which computes the Euclidean distance between the z-score, log normalized higher order parameters of each cell and z-score, log normalized higher order parameter values of the centroid (obtained through k-means clustering) of each cluster. We assigned the cluster as the cluster demonstrating the least Euclidean distance.

### Sankey dataset calculation

To plot the temporal Sankey datasets, we first divided the clusters into pertinent groups: SG1, SG2, SG3, and SG4 for 2.5 hours datasets and CG1, CG2, CG3, and CG4 for 8hrs datasets. We then linked the temporal transition of 2.5hrs trajectories to their respective 8hrs trajectories. Hence, we plotted three temporal segments of 2.5hrs trajectories. The stability fraction is computed as the fraction of cells entering and staying in a short durations class from each of the 4 behavior classes for each 2.5-hr time segment. For instance, SG1 and SG4 had higher stability fractions since most of the cells tended to stay in SG1 or SG4 if they started in SG1 or SG4. However, there were smaller, similar fractions of cells entering SG2 and SG3 from each of the four short duration behavior classes.

### Entropy Calculations

#### Spatial Entropy

To compute spatial entropy, we used the occurrences of 8hrs trajectories of the biological condition under consideration. We then computed entropy using the following equation:

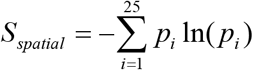

The occurrences are probabilities (*p*_i_) of cells in each n=25 cluster (i=1,2,3…25).

#### State Entropy

To compute state entropy, we calculated the probability of each group SG1, SG2, SG3, SG4 at each time segment, T1, T2, and T3 (equation 15) and CG1, CG2, CG3, and CG4 for the last part of the temporal Sankey (equation 16).

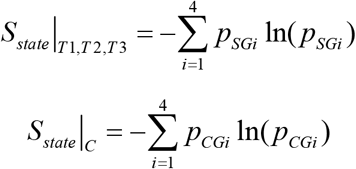

*S*_*state*_ |_*T* 1_, *S*_*state*_ | _*T* 2_, *S*_*state*_|_*T* 3_ are calculated separately from equation (15).

#### Transition Entropy

Transition entropy measures the heterogeneity in the distribution of transition from one-time segment to another.

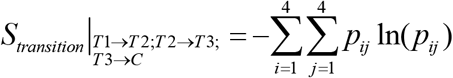

For the computation of *p*_*ij*_ to estimate *S*_*transition*_ |_*T* 1→*T* 2_ ; *S*_*transition*_ |_*T* 2→*T* 3_, we used equation (18) which uses the transition from SGi to SGj (groups of 2.5hrs trajectories). For the calculation of *p*_*ij*_ to estimate *S*_*transition*_| _*T* 3→*C*_, we used equation (19), which uses the transition from SGi (groups of 2.5 hours trajectories) to CGj (groups of 8hrs trajectories).

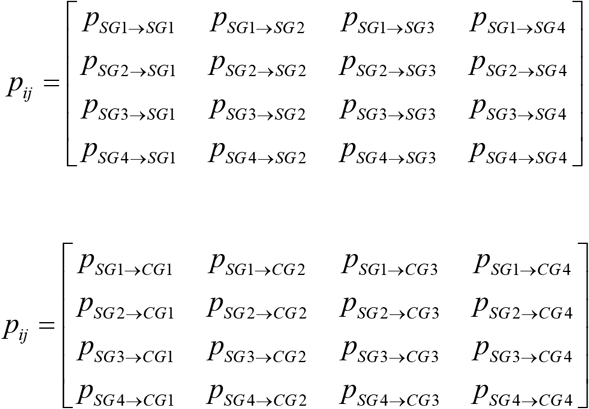

#### Conditional Transition Entropy

We measured the heterogeneity in the distribution in motility groups, given a specific group of a time segment. Three types of conditional transition entropies can link the three-time segments of 2.5hrs trajectories (T1, T2, and T3). *S*_*x* → *i* → *j*_ ; *S*_*i* → *x* → *j*_ ; *S*_*i* → *j* → *x*_ (The temporal transitions are defined as follows-T1: SGx, T2: SGi, T3: SGj, OR, T1: SGi, T2: SGx, T3: SGj, OR, T1: SGi, T2: SGj, T3: SGx, where, SGx is a given group in the respective time segment). 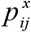 is the probability of transition from SGi to SGj given SGx (at the respective time segment, namely, T1, T2, or T3).

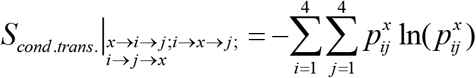

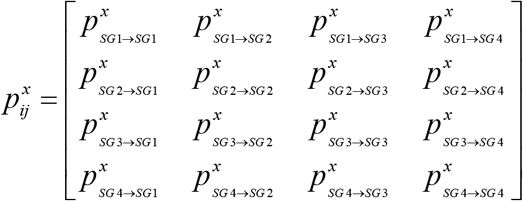

### Predicting 2D vs. 3D

To predict cells moving in either 2D or 3D, we trained a linear SVM for each entropy based on 25 iterations of a 90:10 split of the 60 conditions. The final accuracy results are reported as the average values for each 25 iterations.

## DATA AVAILABILITY

The authors declare that all data supporting the findings in this study are available within the paper and its supplementary information/documents, and on the Phillip lab GitHub repository: https://github.com/PhillipLab-JHU/CaMI

## CODE AVAILABILITY

Detailed descriptions of our approaches and code utilized are provided in the supplementary documents, are available through published literature and on the Phillip lab GitHub repository: https://github.com/PhillipLab-JHU/CaMI

## ACKNOWLEDGEMENTS

We acknowledge financial support for this study from the National Institutes of Health U01AG060903 (JMP, JW), the Johns Hopkins University Older Americans Independence Center of the National Institute on Aging (NIA) under award number P30AG021334 (JMP, JW), and Start-up funds from the Biomedical Engineering Department and the Whiting School of Engineering at Johns Hopkins University.

## AUTHOR CONTRIBUTIONS

JMP conceived the study; JMP, DM, NS conceived data analysis and designed visualization workflows; NZ, HJ, WD, JMP, AJ, BS, LV, AA, DR, AM, LC, and JW performed experiments, collected data, and contributed reagents; DM, JMP, NS, NZ, PK, CM performed formal analysis; JMP, DM, NS, NZ, JW interpreted results; JMP, DM, NS wrote manuscript; all authors contributed to reviewing and editing manuscript; JMP supervised study and secured funding.

## PREPRINT

The manuscript was posted as a preprint on BioRxiv: https://www.biorxiv.org/content/10.1101/2022.09.21.508955v1

