## Supplementary Information for "Profiling Dynamic Patterns of Single-cell Motility"

---

**Supplementary Figure 1. Data and harmonization across multiple datasets. A.** Trajectory of a single ht1080 cell showing the effects of varying interval times (2 to 10 minutes per 30-second steps). Box and whisker plot below show the magnitude of the geometric concordance with varying intervals. **B**. Trajectory of an ht1080 cell at a time resolution of 4 minutes (blue) and the predicted 4-minute intervals based on interpolation (purple) and pseudo-Monte Carlo method (pMC-2.01, brown). Box and whisker plot below show the magnitude of the geometric concordance. **C**. Trajectory of an ht1080 cell at a time resolution of 6 minutes (blue) and the predicted 6-minute intervals based on interpolation (purple) and pseudo-Monte Carlo method (pMC-2.01, brown). Box and whisker plot below show the magnitude of the geometric concordance. In both cases (B and C), the starting time resolution was 8 minutes.

**Supplementary Figure 2. Sensitivity analysis across pMC factors. A.** Mean squared displacements (MSD) of primary human dermal fibroblasts (n=873 cells) as a function of time lag. Data indicates that interpolation and pMC1 exhibits significant deviations relative to original trajectories at short time-scales. **B**. Sensitivity analysis showing comparisons in cell motility parameters for original, interpolation, and four pMC factors.

**Supplementary Figure 3. Characterization of short duration cell motility clusters. A.** Communality factor analysis for the use of eight APRW motility parameters. Bar plots show the average communality score for eight APRW-based parameters relative to other base motility features (Heteromotility, Kimmel et al. PLOS Computational Biology, 2018). **B**. Line plot shows the number of clusters and the inertia/silhouette ratios. **C**. Heatmap of cluster-coherence showing that cells classified within each motility cluster were close to their cluster centroids. The characteristic 'staircase' pattern indicates that cells clustered within each group exhibit short distances to the centroid of the respective clusters. **D**. Radar plot showing the magnitude of the eight motility parameters for the seventeen short duration motility clusters. Magnitude of the motility parameters per cluster highlight some of the fundamental differences among clusters.

**Supplementary Figure 4. Classification accuracy and stability of short-duration motility clusters. A-B.** Confusion matrix showing the classification predictions using linear SVM for cells 90:10 split of data (A) and the prediction classifications using linear SVM of 75,000 simulated cell trajectories (B).

**Supplementary Figure 5. Characterization of long duration cell motility clusters. A.** Line plot shows the number of clusters and the inertia/silhouette ratios. **B**. Heatmap of cluster-coherence showing that cells classified within each motility cluster were close to their cluster centroids. The characteristic 'staircase' pattern indicates that cells clustered within each group exhibit short distances to the centroid of the respective clusters. **C-D**. Mean squared displacements per long duration motility clusters separated based on the four long duration behavior classes (CG1-CG4) (C). On the right we show the probability density function profiles of the angular velocity magnitudes for cells within each of the four long duration behavior classes (D). Data indicates that cells in CG2 and CG3 are more spatially persistent than cells in CG1 and CG4. Low circularity values (more ellipsoidal profiles) indicate behavior classes having high spatial persistence. **E**. Radar plot showing the magnitude of the eight motility parameters for the twenty-five long duration motility clusters.

**Supplementary Figure 6. Classification accuracy and stability of long-duration motility clusters. A-B.** Confusion matrix showing the classification predictions using linear SVM for cells 90:10 split of data (A) and the prediction classifications using linear SVM of 25,000 simulated cell trajectories (B).

**Supplementary Figure 7. Classification of genetic perturbations and drug treatments in LY12 and ht1080 cells. A-L.** Different biological conditions shown on the 2D tSNE maps of the 409 conditions. Conditions include LY12 CucE (A), LY12 Docetaxel (B), LY12 Jasplakinolide (C), LY12 Latrunculin B (D), LY12 LI5 (E), LY12 Taxol (F), LY12 Vincristine (G), LY12 Y27632 (H), ht1080 high and low density in 2D exposed to Reparaxin (I), ht1080 high and low density in 3D exposed to Reparaxin (J), ht1080 cells depleted of WASF3 at high and low cell densities (K), ht1080 cells depleted of ARP2/3 exposed to IL6 inhibitor Tocilizumab (L).

**SUPPLEMENTARY TABLES/DATASETS.**

1. **Supplementary Table 1**. st_XY_condition.csv -- contains time, XY coordinates, and biological condition number for each short-term trajectory. The 'st_id' column maps each trajectory to rows in the higher order parameter dataframe (# 2). Columns are organized follows: ['Biological Condition No.', 'Cell Number', 'Time Point', 'X', 'Y', 'st_id']
2. **Supplementary Table 2**. st_HO_params.csv -- contains raw higher order features, cluster #, sg class, tSNE coordinate, and condition data for each 2.5-hour trajectory. The 'st_id' column maps each row of this data frame to each trajectory in st_XY_condition.csv. Columns are organized as follows: ['MSD10', 'MSD100', 'Pp', 'Pnp', 'Dp', 'Dnp', 'Dtot', 'phi', 'st_id','cluster', 'sg', 'tSNE_x', 'tSNE_y', 'condition', 'condition_name','cell_type']
3. **Supplementary Table 3**. st_HO_params_scaled.csv -- contains Z-score and log-normalized higher order features, cluster #, sg class, tSNE coordinate, and condition data for each 2.5-hour trajectory. The 'st_id' column maps each row of this data frame to each trajectory in st_XY_condition.csv. Columns are organized as follows: ['MSD10', 'MSD100', 'Pp', 'Pnp', 'Dp', 'Dnp', 'Dtot', 'phi', 'st_id','cluster', 'sg', 'tSNE_x', 'tSNE_y', 'condition', 'condition_name','cell_type'].
4. **Supplementary Table 4.** lt_XY_condition.csv -- contains time, XY coordinates, and biological condition number for each 8-hour trajectory. The 'lt_id' column maps each trajectory to rows in the higher order parameter dataframe (# 5). Columns are organized follows: ['Biological Condition No.', 'Cell Number', 'Time Point', 'X', 'Y', 'lt_id']
5. **Supplementary Table 5.** lt_HO_params.csv -- contains raw higher order features, cluster #, cg class, tSNE coordinate, and condition data for each 8-hour trajectory. The 'lt_id' column maps each row of this data frame to each trajectory in lt_XY_condition.csv. Columns are organized as follows: ['MSD10', 'MSD100', 'Pp', 'Pnp', 'Dp', 'Dnp', 'Dtot', 'phi', 'lt_id', 'cluster', 'cg', 'tSNE_x', 'tSNE_y', 'condition', 'condition_name','cell_type']
6. **Supplementary Table 6**. lt_HO_params_scaled.csv -- contains Z-score and log-normalized higher order features, cluster #, cg class, tSNE coordinate, and condition data for each 8-hour trajectory. The 'lt_id' column maps each row of this data frame to each trajectory in lt_XY_condition.csv. Columns are organized as follows: ['MSD10', 'MSD100', 'Pp', 'Pnp', 'Dp', 'Dnp', 'Dtot', 'phi', 'lt_id', 'cluster', 'cg', 'tSNE_x', 'tSNE_y', 'condition', 'condition_name','cell_type']

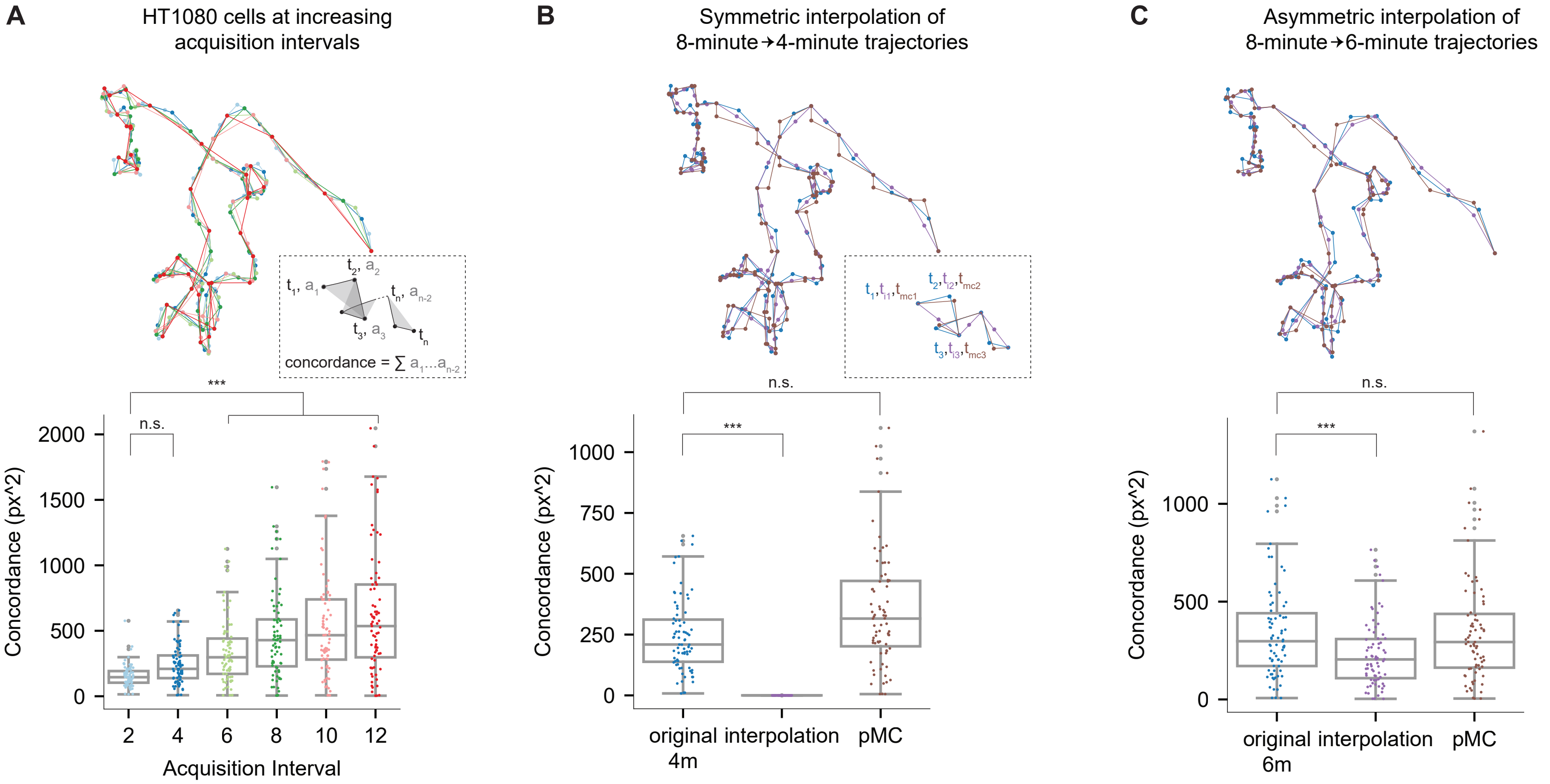


Supplementary Figure 1

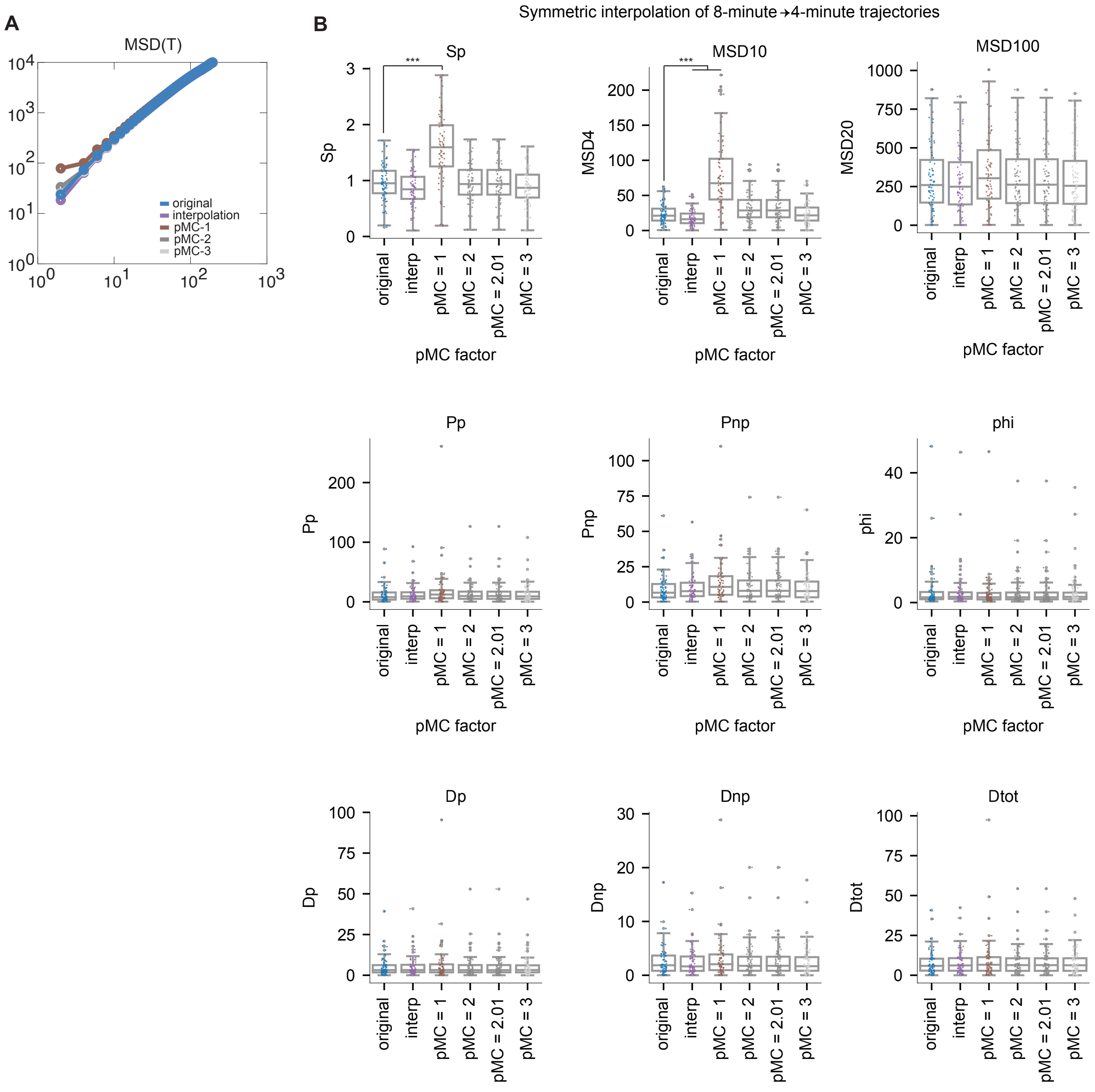


Supplementary Figure 2

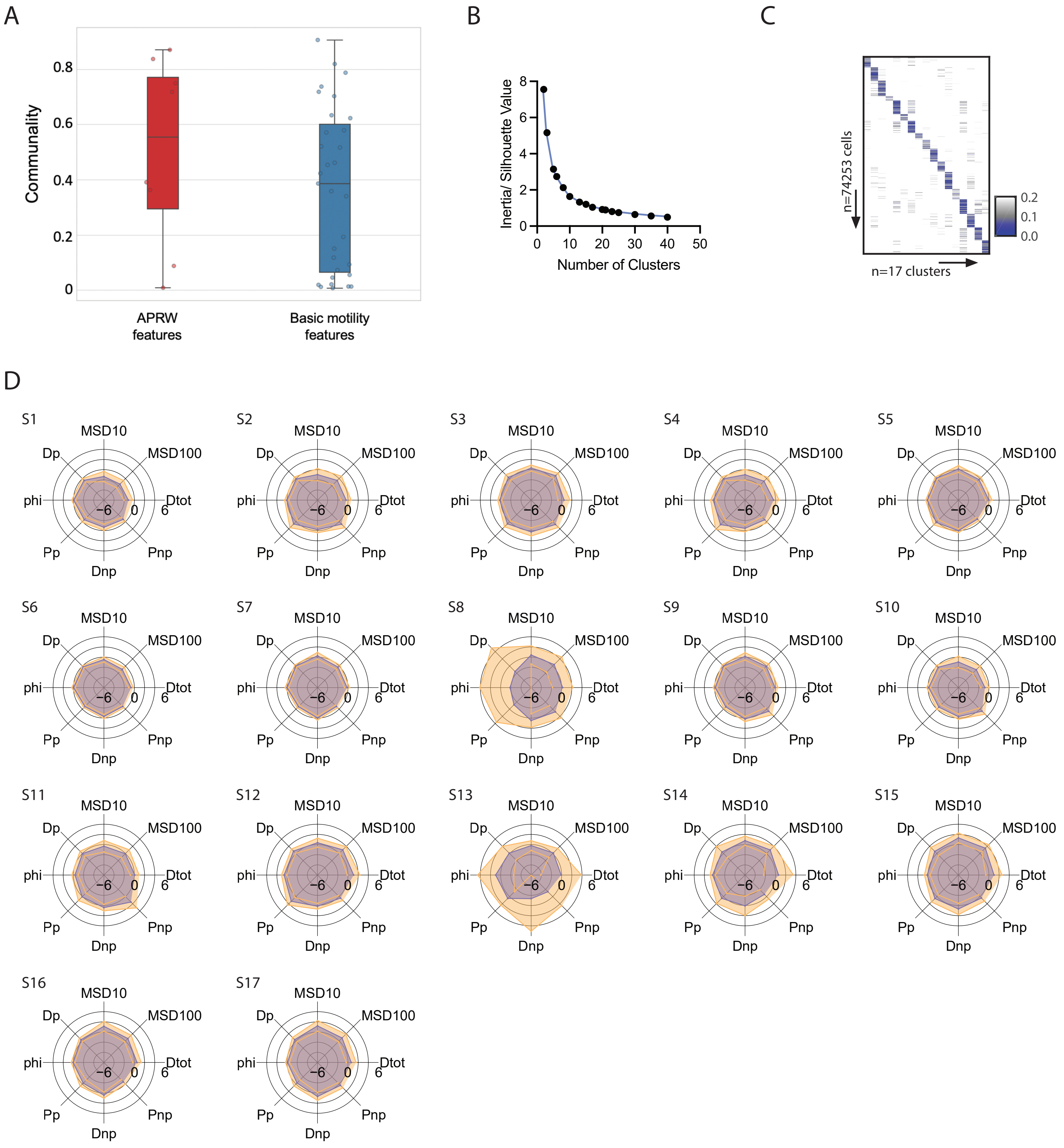


Supplementary Figure 3

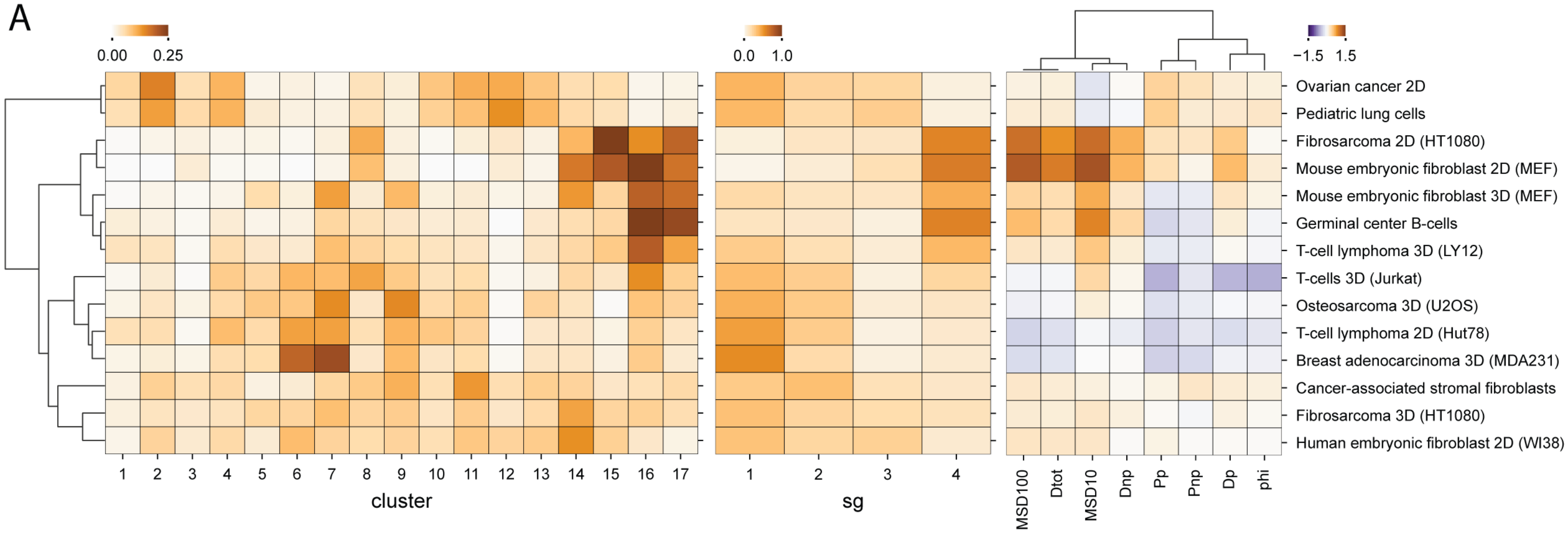


T-cell Lymphoma motility in response to Actin perturbation

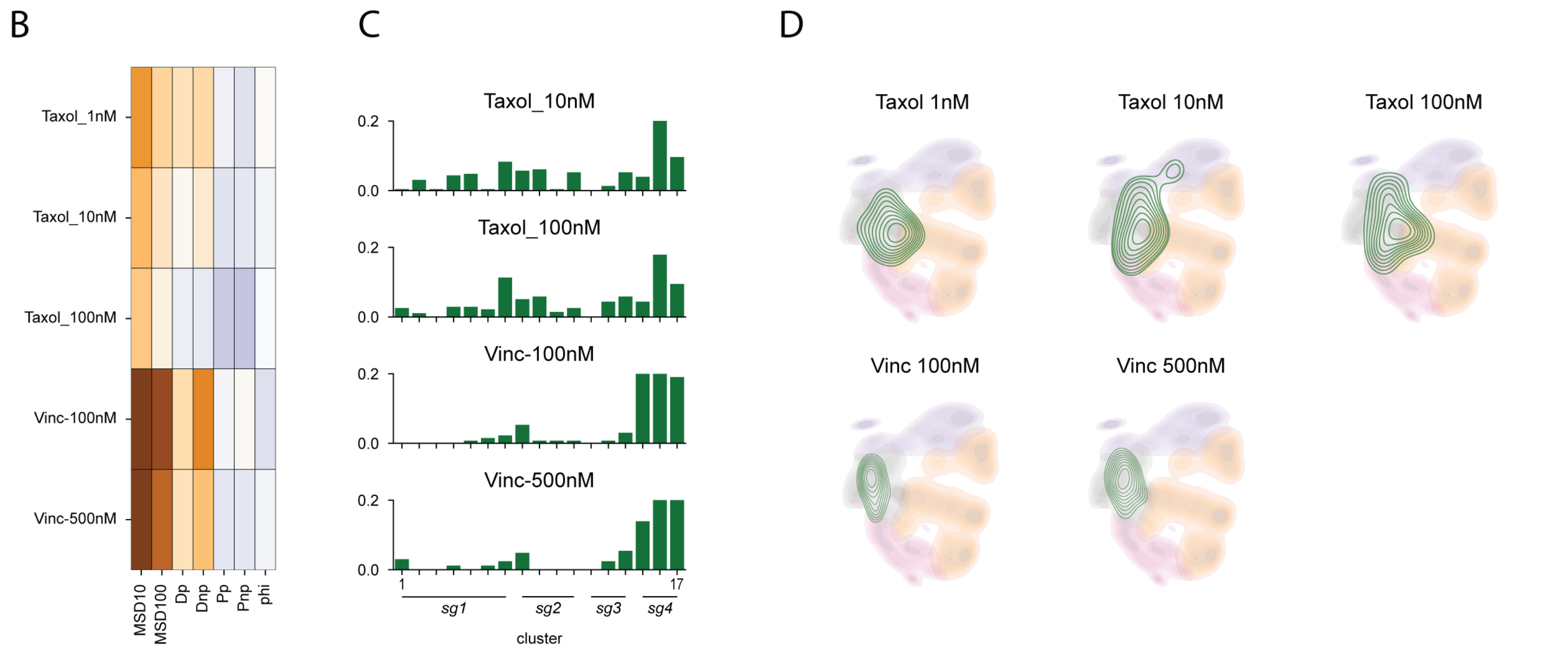


Supplementary Figure 4

A

90:10 (Training:Validation)

| | 1 | 2 | 3 | 4 | 5 | 6 | 7 | 8 | 9 | 10 | 11 | 12 | 13 | 14 | 15 | 16 | 17 |
|---|---|---|---|---|---|---|---|---|---|---|---|---|---|---|---|---|---|
| **1** | 99 | | | 2.3 | | | | | 1.1 | 0.8 | | | | | | | |
| **2** | | 99.6 | | | | | | 0.8 | | | | | | | | | |
| **3** | | 0.4 | 99.3 | 1.1 | | | | 1.7 | | | | 0.6 | | | | | |
| **4** | 0.5 | | | 94.3 | | 0.6 | | | | | | | | | | | |
| **5** | | | | 1.1 | 99.6 | | | | | | | | | 0.7 | | | |
| **6** | | | | | | 97.7 | | | | | | 0.3 | | | | | |
| **7** | | | | | | | 99.6 | | 0.5 | | | | | | | | |
| **8** | | | | | | | | 96.7 | | | | | 1.0 | | | | |
| **9** | 0.5 | | | | | | | | 97.8 | | | | | | | | |
| **10** | | | | 1.1 | | | | | | 98.3 | | | | | | | |
| **11** | | | | | | | | | | | 100 | | | | | | |
| **12** | | | | | | 0.8 | | | | | | 99.1 | | | | | |
| **13** | | | 0.7 | | | 0.8 | | | | | | | 99 | | | | |
| **14** | | | | | | | | 0.8 | | | | | | 99.3 | | 0.8 | |
| **15** | | | | | | | | | | 0.8 | | | | | 100 | | |
| **16** | | | | | | | 0.4 | | | | | | | | | 99.2 | |
| **17** | | | | | 0.4 | | | | | | | | | | | | 100 |

Accuracy = 98.7%%

| | | | | | | | | | | | | | | | | | |
|---|---|---|---|---|---|---|---|---|---|---|---|---|---|---|---|---|---|
| **PP V** | 99 | 99.6 | 99.3 | 94.3 | 99.6 | 97.7 | 99.6 | 96.7 | 97.8 | 98.3 | 100 | 99.1 | 99 | 99.3 | 100 | 99.2 | 100 |
| **FDR** | 1.0 | 0.4 | 0.7 | 5.7 | 0.4 | 2.3 | 0.4 | 3.3 | 2.2 | 1.7 | 0.0 | 0.9 | 1.0 | 0.7 | 0.0 | 0.8 | 0.0 |

B

75,000 simulated cells

| | 1 | 2 | 3 | 4 | 5 | 6 | 7 | 8 | 9 | 10 | 11 | 12 | 13 | 14 | 15 | 16 | 17 |
|---|---|---|---|---|---|---|---|---|---|---|---|---|---|---|---|---|---|
| **1** | 97.9 | | | 1.37 | 0.24 | | 0.26 | | 0.32 | | | | | | | | |
| **2** | | 97.9 | 0.17 | | | | | 0.08 | | | 0.75 | 0.06 | 0.06 | | 0.04 | | |
| **3** | | 0.6 | 98.2 | | | 0.14 | | 0.36 | | | | 0.23 | 0.23 | | | | |
| **4** | 0.24 | | | 81 | 0.24 | 0.06 | | | | 0.18 | | 0.06 | 0.06 | | | | 0.06 |
| **5** | 0.08 | | | 1.69 | 98.5 | 0.24 | 0.11 | 0.03 | | | | | | 0.68 | | | 0.03 |
| **6** | | | 1.4 | 1.28 | 0.12 | 96.5 | | | | | | 0.28 | 0.96 | | | | 1.4 |
| **7** | 0.35 | | | 0.02 | 0.15 | | 98.8 | 0.05 | 0.85 | | | | | 0.55 | | 0.4 | |
| **8** | 0.04 | | 0.41 | 0.08 | | 0.21 | | 95.7 | | | 0.04 | | 0.9 | 0.25 | | 0.04 | 0.62 |
| **9** | 1.46 | | | 0.07 | | 0.02 | 0.41 | | 96.8 | 1.97 | 0.07 | | | | 0.58 | 0.02 | |
| **10** | 0.46 | | | 1.02 | | | | 0.02 | 0.5 | 98.2 | 0.02 | 0.15 | | | 0.45 | | |
| **11** | | 0.56 | | | | | | 0.52 | 0.04 | | 96.9 | | 0.32 | | | 0.16 | |
| **12** | | 0.24 | 0.25 | 1.0 | | 0.18 | | 0.01 | | 0.15 | | 99.4 | | | 0.42 | | |
| **13** | | | 1.1 | | | 0.22 | | 0.22 | | | 0.02 | | 98.1 | | | | 0.9 |
| **14** | | | | | 0.2 | | 0.22 | 0.22 | | | | | | 96.1 | | 0.37 | 0.22 |
| **15** | | 0.51 | | 0.03 | | | | | 0.77 | 0.34 | 0.15 | 0.07 | 0.02 | | 98.5 | | |
| **16** | | | | | 0.04 | | 0.11 | 0.6 | 0.04 | | 1.02 | | 0.19 | 1.66 | | 98.5 | 0.04 |
| **17** | | | | 0.06 | 0.37 | 0.73 | | 0.43 | | | | | 0.27 | 1.18 | | 0.06 | 97.9 |

Accuracy = 96.7%

| | | | | | | | | | | | | | | | | | |
|---|---|---|---|---|---|---|---|---|---|---|---|---|---|---|---|---|---|
| **PP V** | 97.9 | 97.9 | 98.2 | 81 | 98.5 | 96.5 | 98.8 | 95.7 | 96.8 | 98.2 | 96.9 | 99.4 | 98.1 | 96.1 | 98.5 | 98.5 | 97.9 |
| **FDR** | 2.1 | 2.1 | 1.8 | 19.0 | 1.5 | 3.5 | 1.2 | 4.3 | 3.2 | 1.8 | 3.1 | 0.6 | 1.9 | 3.9 | 1.5 | 1.5 | 2.1 |

Supplementary Figure 4

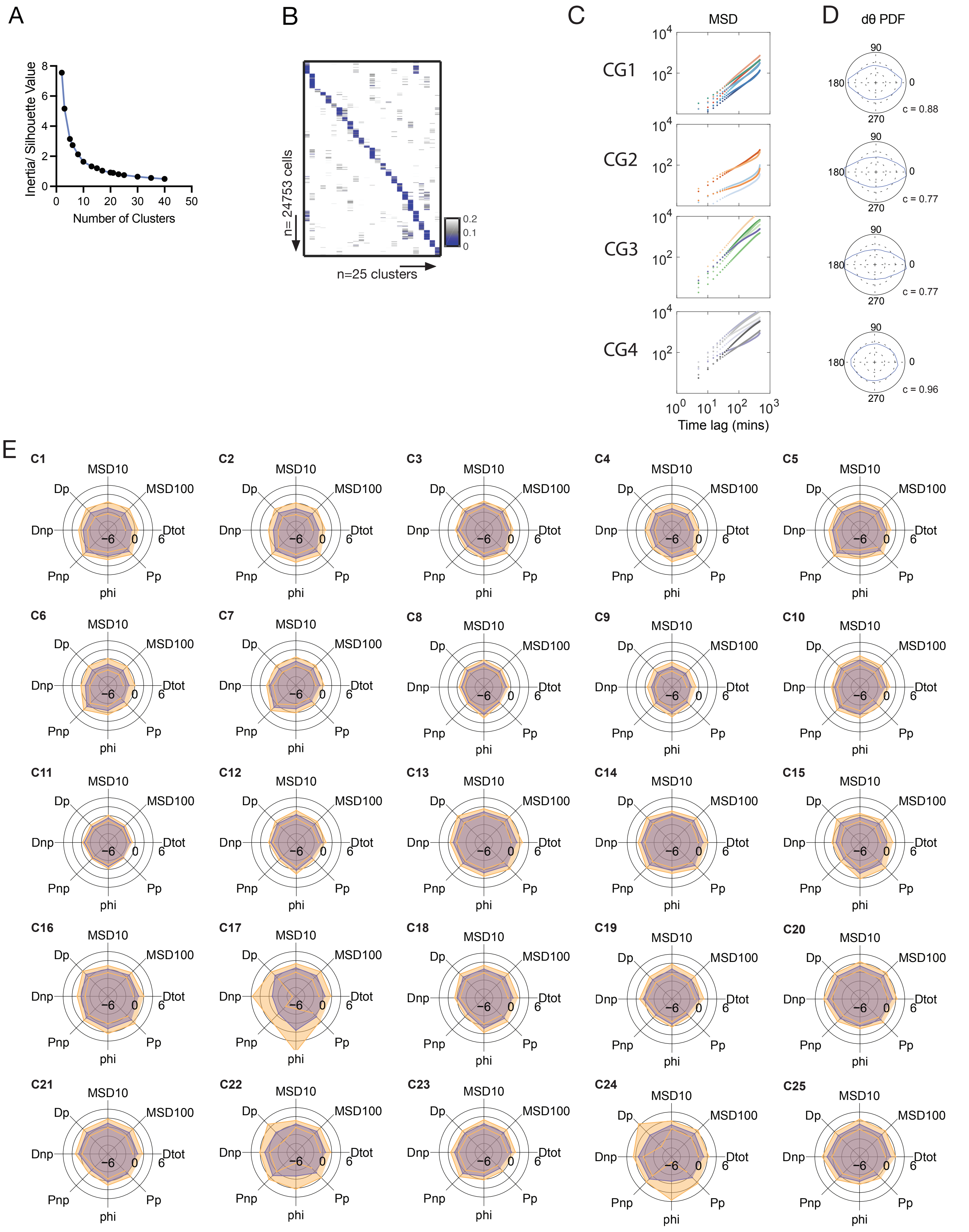


Supplementary Figure 5

A

### 90:10 (Training:Validation)

| | 1 | 2 | 3 | 4 | 5 | 6 | 7 | 8 | 9 | 10 | 11 | 12 | 13 | 14 | 15 | 16 | 17 | 18 | 19 | 20 | 21 | 22 | 23 | 24 | 25 |
|---|---|---|---|---|---|---|---|---|---|---|---|---|---|---|---|---|---|---|---|---|---|---|---|---|---|
| **1** | 96.3 | | | | | | | | | 1.8 | | | | | | | | | | | | | | | |
| **2** | | 96.9 | | | | | | | | | | | | 3.7 | | | | | | | | | | | |
| **3** | | | 91.7 | | | | | | | | | | | | | | | | | | | | | | |
| **4** | | | 4.2 | 100 | | | | | | | | | | | | | | | | | | | | | |
| **5** | | | | | 96.0 | | | 3.7 | | | | | | | | | | | | | | | | | |
| **6** | | | | | | 96.4 | | | | | | | | | | | | | | | | | | | |
| **7** | | | | | | | 98.1 | | | | | | | | | | | 3.5 | | | | | | | |
| **8** | | | 4.2 | | | | 1.9 | 93.6 | | | | 9.5 | 1.5 | | 2.3 | | | 1.8 | | | | | | | |
| **9** | | | | | | | | | 98.1 | | | | | | | | | | | | | | | | |
| **10** | | | | | | 3.6 | | | | 96.4 | | | | | | | | | | | | | | | |
| **11** | | | | | | | | | 1.9 | | 100 | | | | | | | | | | | | | | |
| **12** | | | | | | | | | | | | 90.5 | | | | | 2.2 | | | | | | 1.6 | | |
| **13** | | 1.0 | | | | | | 3.7 | | | | | 98.5 | | | | | | | | | | | | |
| **14** | | 1.0 | | | 2.0 | | | | | | | | | 96.3 | | | | | | | | 1.7 | | | |
| **15** | 1.3 | | | | | | | | | | | | | | 95.5 | | | | | | | | | | |
| **16** | | | | | 2.0 | | | | | | | | | | | 100 | | | | | | | | | |
| **17** | 1.3 | | | | | | | | | | | | | | | | 97.8 | | | | | | | | |
| **18** | | | | | | | | | | | | | | | | | | 94.7 | | | | | | | |
| **19** | | | | | | | | | | | | | | | | | | | 100 | | | 1.7 | | | |
| **20** | | | | | | | | | | | | | | | | | | | | 100 | | | | | |
| **21** | 1.3 | | | | | | | | | | | | | | | | | | | | 100 | | | | 3.4 |
| **22** | | | | | | | | | | | | | | | | | | | | | | 96.6 | | | |
| **23** | | 1.0 | | | | | | | | | | | | | | | | | | | | | 98.4 | | |
| **24** | | | | | | | | | | 1.8 | | | | | | | | | | | | | | 100 | |
| **25** | | | | | | | | | | | | | | | 2.3 | | | | | | | | | | 96.6 |

| | 1 | 2 | 3 | 4 | 5 | 6 | 7 | 8 | 9 | 10 | 11 | 12 | 13 | 14 | 15 | 16 | 17 | 18 | 19 | 20 | 21 | 22 | 23 | 24 | 25 |
|---|---|---|---|---|---|---|---|---|---|---|---|---|---|---|---|---|---|---|---|---|---|---|---|---|---|
| **PPV** | 96.3 | 96.9 | 91.7 | 100 | 96.0 | 96.4 | 98.1 | 93.6 | 98.1 | 96.4 | 100 | 90.5 | 98.5 | 96.3 | 95.5 | 100 | 97.8 | 94.7 | 100 | 100 | 100 | 96.6 | 98.4 | 100 | 96.6 |
| **FDR** | 3.7 | 3.1 | 8.3 | 0.0 | 4.0 | 3.6 | 1.9 | 6.4 | 1.9 | 3.6 | 0.0 | 9.5 | 1.5 | 3.7 | 4.5 | 0.0 | 2.2 | 5.3 | 0.0 | 0.0 | 0.0 | 3.4 | 1.6 | 0.0 | 3.4 |

Accuracy = 97.2

B

### 25,000 simulated cells

| | 1 | 2 | 3 | 4 | 5 | 6 | 7 | 8 | 9 | 10 | 11 | 12 | 13 | 14 | 15 | 16 | 17 | 18 | 19 | 20 | 21 | 22 | 23 | 24 | 25 |
|---|---|---|---|---|---|---|---|---|---|---|---|---|---|---|---|---|---|---|---|---|---|---|---|---|---|
| **1** | 98.50 | | | 0.11 | | 0.50 | | | | 0.11 | | | | | 0.11 | | 0.11 | | | | 1.57 | | | | 0.34 |
| **2** | | 97.70 | | | | | | | | | 0.10 | 0.19 | 0.10 | 0.19 | | | | | 0.68 | | | 0.29 | 1.36 | | |
| **3** | | | 92.10 | 1.53 | | 0.22 | 1.53 | 0.44 | | | | 0.22 | | | 0.87 | | | 0.87 | 1.53 | | | 0.44 | | | |
| **4** | | | 0.16 | 94.40 | | 0.48 | 2.38 | | | | | | | 0.16 | | | | 1.27 | | 0.16 | | | | | 1.27 |
| **5** | | | | | 96.50 | | | 0.19 | | | | | 0.56 | 2.11 | | 0.58 | | | | | | | | | |
| **6** | 0.82 | | 1.43 | 0.41 | | 88.40 | | | 2.04 | 1.63 | | | | | | | | | | | | | | | |
| **7** | | | | 0.51 | 0.07 | | 90.90 | 0.37 | | | | | | | 0.15 | 0.22 | | 0.07 | | | | | | | 0.29 |
| **8** | | | | | 0.64 | | 6.22 | 95.50 | | | | 0.64 | 0.48 | | 0.32 | 0.32 | | 1.59 | | | | | | | |
| **9** | | | | | | 1.12 | | | 97.00 | 0.78 | 0.69 | 0.09 | | | | | | | | 1.12 | | | | 1.29 | |
| **10** | 1.02 | | | | | 0.66 | | | 0.36 | 98.40 | | | | | | | 0.87 | | | | | | | 0.44 | |
| **11** | | | | | | | | | 0.51 | | 97.20 | 0.93 | | 0.17 | | | | | | 0.59 | | | 1.53 | 0.42 | |
| **12** | | | | | 0.59 | | | 0.44 | | | 1.18 | 93.90 | 0.15 | | | 0.30 | 3.55 | | | | | | | 2.36 | |
| **13** | | 0.78 | | | 0.65 | | | 0.91 | | | | 0.33 | 97.80 | 0.65 | | | | 0.39 | | | | 0.13 | | | |
| **14** | | 0.37 | | | | | | | 0.37 | | | | 0.75 | 90.60 | | | 0.37 | | | 0.37 | | | | 0.75 | |
| **15** | 0.11 | | 0.33 | 0.22 | | 0.76 | 4.26 | 0.22 | | | | 0.13 | | | 97.20 | 0.11 | 0.22 | | | | 3.06 | | | | 1.64 |
| **16** | 0.18 | | | 0.18 | 0.71 | | 0.35 | | | | | 0.53 | | 0.35 | | 97.20 | 0.35 | | | | 3.70 | | | | 0.71 |
| **17** | 0.33 | | | | | | | | | 0.08 | | 0.66 | | | | 0.25 | 95.90 | | | | 0.25 | | | 0.33 | |
| **18** | | | 0.27 | 0.09 | | | 0.89 | 0.63 | | | | | 0.18 | | | | | 96.90 | | | | | | | |
| **19** | | 0.62 | 1.04 | 0.21 | | | | | | | | | | | | | | | 96.70 | | | 0.21 | 0.94 | | |
| **20** | | | 0.53 | | | 1.24 | | | 0.62 | | 0.18 | | | | | | | | 0.44 | 97.40 | | | 0.79 | | |
| **21** | 0.06 | | | | | | | | | | | 0.26 | | | 0.66 | 0.12 | 0.18 | | | | 93.80 | | | | 0.12 |
| **22** | | 0.72 | 0.54 | 0.27 | | | | | | | | | 0.45 | | | | | 0.81 | 0.63 | | | 99.30 | | | |
| **23** | | 0.26 | | | | | | | | | 1.02 | | | 0.17 | | | | | 0.51 | 0.68 | | | 95.70 | | |
| **24** | | | | | | | | | 0.43 | 0.17 | 0.17 | 0.52 | | | | | | | | | | | | 95.80 | |
| **25** | 0.07 | | | 0.57 | | 0.07 | 0.85 | | | | | | | | 0.35 | | | | | | 1.70 | | | | 97.20 |

| | 1 | 2 | 3 | 4 | 5 | 6 | 7 | 8 | 9 | 10 | 11 | 12 | 13 | 14 | 15 | 16 | 17 | 18 | 19 | 20 | 21 | 22 | 23 | 24 | 25 |
|---|---|---|---|---|---|---|---|---|---|---|---|---|---|---|---|---|---|---|---|---|---|---|---|---|---|
| **PPV** | 98.5 | 97.7 | 92.1 | 94.4 | 96.5 | 88.4 | 90.9 | 95.5 | 97.0 | 98.4 | 97.2 | 93.9 | 97.8 | 90.6 | 97.2 | 97.2 | 95.9 | 96.9 | 96.7 | 97.4 | 93.8 | 99.3 | 95.7 | 95.8 | 97.2 |
| **FDR** | 1.5 | 2.3 | 7.9 | 5.6 | 3.5 | 11.6 | 9.1 | 4.5 | 3.0 | 1.6 | 2.8 | 6.1 | 2.2 | 9.4 | 2.8 | 2.8 | 4.1 | 3.1 | 3.3 | 2.6 | 6.2 | 0.7 | 4.3 | 4.2 | 2.8 |

Accuracy = 95.6%

Supplementary Figure 6

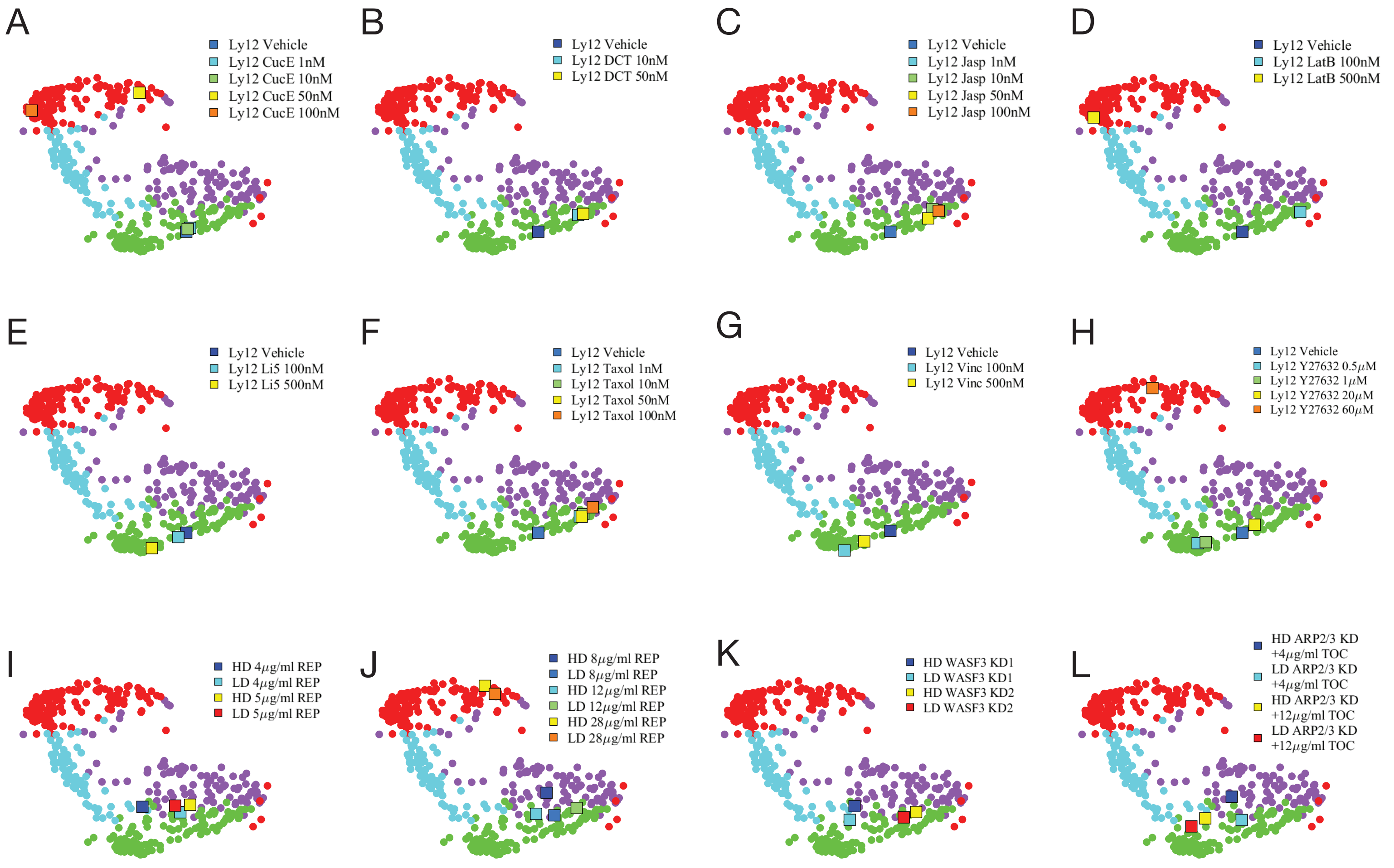


Supplementary Figure 7
